# Functional promiscuity of small multidrug resistance transporters from *Staphylococcus aureus*, *Pseudomonas aeruginosa*, and *Francisella tularensis*

**DOI:** 10.1101/2022.11.22.517605

**Authors:** Peyton J. Spreacker, Colin J. Porter, Andrea Wegrzynowicz, Will F. Beeninga, Sydnye Demas, Emma N. Powers, Katherine A. Henzler-Wildman

**Affiliations:** Department of Biochemistry, University of Wisconsin-Madison, Madison, WI

**Author notes:** Address correspondence to Katherine A. Henzler-Wildman. Mayo Medical Laboratories, Rochester, MN Running Header: Bacterial multidrug efflux pump promiscuity.

## Abstract

Small multidrug resistance (SMR) transporters efflux toxic substrates from bacterial cells and were recently divided into two subfamilies: specific toxic metabolite transporters and promiscuous drug exporters. These drug exporters are thought to function similarly to EmrE, the model system for this subfamily of SMR transporters. Studies of EmrE homologs indicate that they are able to confer resistance to EmrE substrates in *E. coli* and in their native organisms. Recent work from our lab showed that functional EmrE can confer resistance or susceptibility *in vivo* depending on the drug substrate. Here, we test whether this functional promiscuity of EmrE extends to SMR transporters from three additional human or animal pathogens: SAsmr from *Staphylococcus aureus*, PAsmr from *Pseudomonas aeruginosa*, and FTsmr from *Francisella tularensis*. We find that these SMR homologs can confer either resistance or susceptibility to different toxic substrates in *E. coli*. This demonstrates that the ability of a single transporter to lead to opposite biological outcomes when transporting different substrates is a general property of the promiscuous multidrug transporters in the SMR family. It also suggests the potential for novel antibiotic development targeting these transporters with small molecules that trigger susceptibility. Such a strategy does not require that the target be the primary mode for antibiotic resistance because the goal is not simple inhibition of activity, but rather activation of an alternative transport function that is detrimental to bacteria.

## INTRODUCTION

Multidrug resistance (MDR) transporters actively efflux antibiotics, antiseptics, and other toxic compounds, reducing intracellular concentrations sufficiently to enable bacterial survival. The small multidrug resistance (SMR) transporters are the smallest known active transporters and are found only in bacteria (1). Several studies have implicated SMR transporters in a variety of bacterial processes in addition to toxin efflux including biofilm production (2–4), osmotic stress regulation (5, 6), toxin transport (1, 7, 8), and antibiotic resistance (2, 9, 10). Studying the role of these efflux proteins in antibiotic resistance has become paramount with increased prevalence of SMR transporters in multidrug resistant strains of bacteria (11), but limited understanding of the functional role of the SMR transporters in these strains.

Members of the SMR family are readily identified through sequence analysis (12–14) and have been divided into two subclasses: EmrE-like small multidrug pumps with broad substrate specificity and GdX (formerly SugE) -like transporters that have narrower substrate specificity profile and confer resistance to a smaller set of toxic cations. Recently, the GdX-like subfamily was reclassified as toxic metabolite exporters (13, 15) as a result of the discovery that expression of this subfamily is regulated by a novel family of riboswitches (14, 16, 17) in response to guanidinium^+^. Furthermore, transport assays confirmed efficient 2H^+^/1 guanidinium^+^ antiport and selectivity for guanidinium over other closely related compounds by GdX (13). However, there may be exceptions to the definition of the GdX subfamily as more selective, tightly coupled metabolite exporters and the EmrE subfamily as highly promiscuous exporters of exogenous toxins. For example, MdtIJ, a heterodimer classified as EmrE-like via sequence, was shown to transport spermidine, a metabolite, rather than acting as a multidrug transporter (18, 19). This raises the question of how much functional variation exists within each subfamily, and how well functional phenotypes correlate with bioinformatic classifications, since experimental assessment of transport or drug resistance phenotypes has only been performed for a relatively small number of SMR homologs and limited number of substrates (13, 20–26).

EmrE was the first SMR transporter to be functionally characterized (27), both *in vitro* and *in vivo,* and is still the most extensively studied SMR homolog. EmrE functions as an antiparallel homodimer and is a highly promiscuous transporter that exports polyaromatic cations and quaternary ammonium cations across the inner membrane of *E. coli* (2, 21, 27– 29). The well-established antiport of 2 H^+^/1 substrate by EmrE enables active efflux of substrate by coupling it to inward movement of protons down the proton motive force (27, 30–33). Two glutamate residues (E14_A_ and E14_B_), one from each chain of the homodimer, are required for this activity. The original mechanistic model of coupled 2 H^+^: 1 toxin antiport, was based on early experimental data showing electrogenic transport of a +1 toxin and electroneutral transport of a +2 toxin (31). More recent mechanistic studies have demonstrated that EmrE is not a strictly coupled proton/drug antiporter in the manner of the GdX-subfamily and may be able to perform uniport and symport (32, 34) in addition to antiport. This new free exchange model of EmrE transport has major biological implications, since any of the alternative transport modes should lead to enhanced *susceptibility*, rather than resistance to the small molecule substrate (34).

EmrE has a broad spectrum of substrates, including quaternary ammonium compounds, polyaromatic cations, dyes, and planar toxins, but prior functional screens focused on antibiotics and common multidrug (MDR) efflux pump substrates to define the specificity profile of EmrE relative to other MDR efflux pumps (20, 21, 35, 36). Recently we performed a Biolog phenotypic microarray screen, which includes a much broader array of compounds commonly used for phenotypic screening in bacteria. This revealed that WT-EmrE can indeed confer resistance or susceptibility depending on the small molecule substrate being transported (34), as predicted by the free-exchange model for EmrE transport (32). Here we investigate whether this functional promiscuity extends beyond EmrE to other members of the SMR transporter subfamily.

SMR transporters with high sequence similarity to EmrE can be found throughout the bacterial kingdom and have been classified into subfamilies on the basis of both transporter sequence and their regulatory elements (1). Here we focus on the EmrE-like subfamily proposed to be promiscuous drug exporters rather than the more selective metabolite exporters in the Gdx-like subfamily. There is minimal biochemical characterization of SMR substrate specificity, transport activity or transport mechanism beyond the few well-studied modeled systems of EmrE and Gdx, and even less data on SMR function in the native organism. SAsmr (QacC) from *Staphylococcus aureus* has been shown to confer resistance to ethidium bromide in both *Staphylococcus* and *E. coli* (37). PAsmr from *Pseudomonas aeruginosa* was found to confer resistance to aminoglycosides and dyes in *P. aeruginosa* (38) and PAsmr can transport EmrE substrates *in vitro* (39). One study demonstrated that FTsmr in *Francisella tularensis* is part of the antibiotic resistance response in *F. tularensis* (40). More recently, YnfA from *Shigella flexneri*, was shown to confer resistance to EmrE substrates in the native organism (41). These studies show that EmrE homologs in other bacterial species can confer resistance to similar substrates, but the scope of the assays was limited to common multidrug resistance substrates and detection of resistance phenotypes. Here we analyze SMRs from three bacterial species of interest for the potential to cause disease in humans or animals for comparison with EmrE: *Staphylococcus aureus* (SAsmr) (42–45), *Pseudomonas aeruginosa* (PAsmr) (46, 47), and *Francisella tularensis* (FTsmr) (48). These particular transporters were selected on the basis of disease relevance, sequence diversity (Fig. S1), and availability of prior data. Although the SMR transporter family contains both homodimers and heterodimers, we have selected homodimers for their work both for ease of orthologous expression and because the homodimers have been more widely studied.

To test whether these SMRs can confer resistance *and* susceptibility *in vivo*, we performed an unbiased Biolog phenotypic microarray to profile the metabolic activity of *E. coli* expressing either WT or a non-functional point mutant of each SMR transporter in the presence of 240 different compounds. This screen was performed using heterologous expression in MG1655 Δ*emrE E. coli* to facilitate comparison of substrate profiles and phenotypes of the transporters themselves, without the confounding factor of different intrinsic susceptibility or resistance of *E. coli*, *P. aeruginosa*, *S. aureus*, or *F. tularensis* to the various compounds in the Biolog assay. This experimentally demonstrates that promiscuity in both substrate and mechanism applies more broadly to the EmrE-like SMR subfamily, including SMR homologs from clinical pathogens.

## RESULTS

To compare the substrate specificity profile and functional promiscuity of SAsmr, PAsmr, and FTsmr with EmrE, each transporter was heterologously expressed in MG1655 Δ*emrE E. coli*. Mutation of the critical glutamate residue to glutamine (E14Q-EmrE, E14Q- PAsmr, E13Q-SAsmr, E13Q-FTsmr) blocks transport and is a well-established non-functional mutant of SMR transporters. Both WT and non-functional mutants of each SMR homolog express equally well and are inserted into the membrane with similar efficiency *in E. coli* (Fig. S2D, E), and there is minimal difference in growth of *E. coli* expressing each transporter (Fig. S2A-C).

### SMR transporters have broad substrate profiles

To broaden our understanding of the substrate specificity profiles for these transporters and to assess whether they only confer resistance or can confer both resistance and susceptibility in a substrate-specific manner, we performed Biolog Functional Phenotyping Microarrays using the chemical sensitivity panel (49). This panel of ten 96-well microplates with 240 different compounds includes many drug-like compounds of interest for understanding SMR activity. Many known SMR substrates are polyaromatic cations, and this commercially available assay is designed to avoid interference from the natural fluorescence of such compounds, enabling a single consistent assay format for characterization across several hundred potential substrates selected without bias towards common MDR efflux pump substrates. Our recent Biolog phenotypic microarray screen of *E. coli* revealed that EmrE confers resistance or susceptibility depending on the identity of the substrate, an unprecedented result (34). Here we repeat this screen on SAsmr, PAsmr, and FTsmr expressed in MG1655 Δ*emrE E. coli* cells.

The Biolog screen was ran in parallel with *E. coli* expressing either wildtype or non- functional SMR transporter and measures NADH production over 24 hours using a colorimetric indicator. The area under the curve represents the total metabolic activity and the difference between cells expressing functional and non-functional transporter (ΔArea) reflects the impact of transporter activity on NADH production. If NADH production is greater when wildtype transporter was expressed, it indicates that transport activity is beneficial to cell growth and metabolism and the SMR transporter confers resistance to that compound. If NADH production was greater when the non-functional point mutant of the same transporter was expressed, it indicates that SMR activity is detrimental and the SMR transporter confers susceptibility to that compound (see methods for selection criteria).

The entire assay comparing the impact of functional vs non-functional transporter for each SMR homolog as ran in duplicate. The correlation between total metabolic output for the two replicates under each condition (WT or non-functional transporter), was high (Fig. S3). For each screen, ΔArea was calculated for individual wells and scored as positive (resistance) or negative (susceptibility) hits if the difference was above or below the 10% trimmed mean, as described in the methods. Each set of Biolog plates contained four wells at two different concentrations of a single compound. Since the entire assay was ran in duplicate for each transporter, this results in a total of 8 wells for a single compound, or a maximum possible score of ±8. To account for the non-zero rate of false-positives or false-negatives in scoring individual wells, as well as the potential that the transporter could confer resistance or susceptibility at one compound concentration but not the second concentration in the Biolog plates, we selected any compound that scored ≥ +3 as a resistance hit or ≤ -3 as a susceptibility hit. All the known EmrE resistance substrates (methyl viologen and acriflavine) that are present in the Biolog compound set are selected as resistance hits using this scoring system, confirming the validity of this scoring method. The screen identified compounds in both categories, resistance, and susceptibility, for all four of the SMR homologs and generated a large list of potential hits (Fig. 1, S4, Table S1).

**Figure 1.**
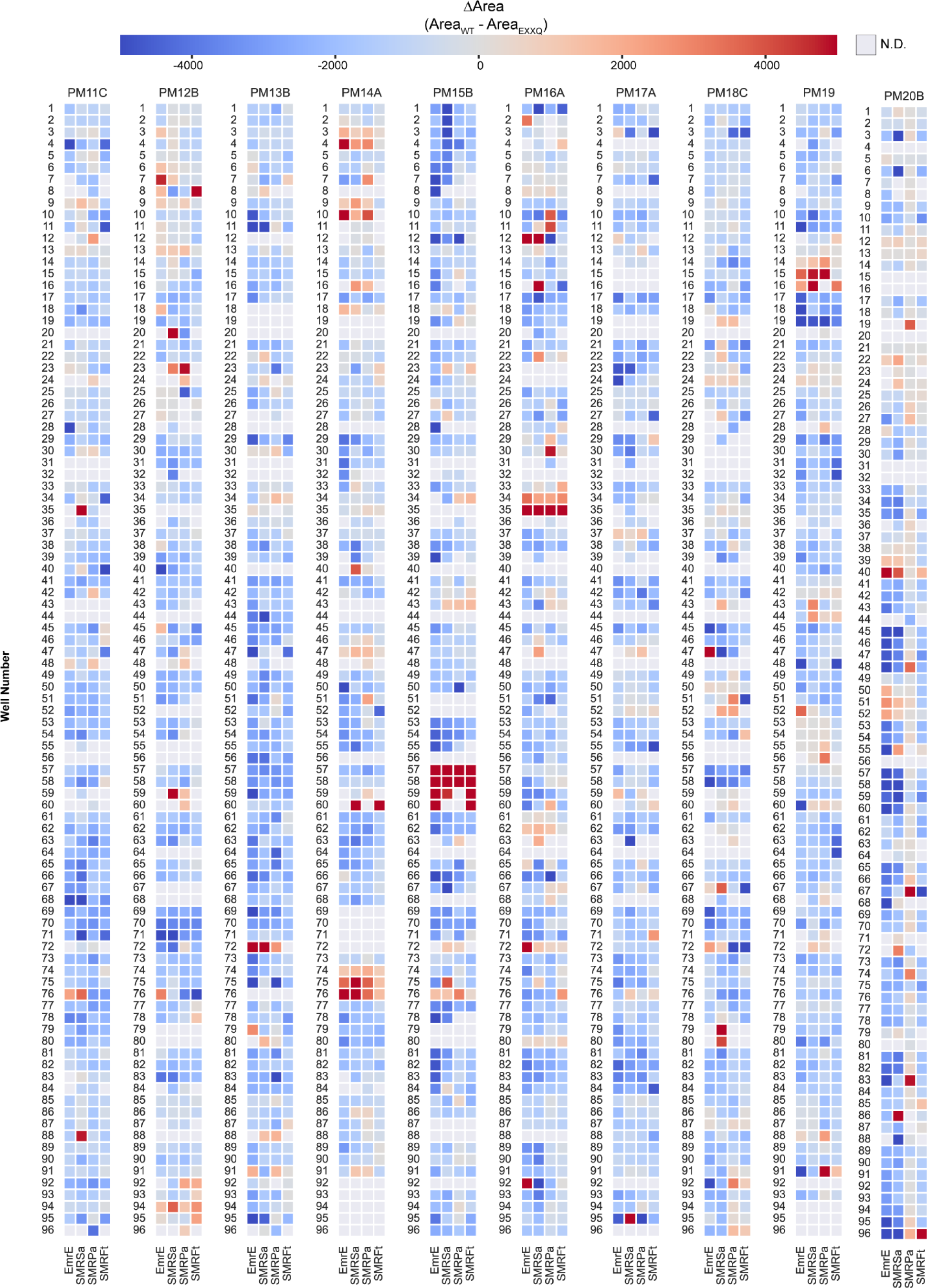
(previous page): Biolog microarray data for all plates. Heat maps of the small molecule plates from the Biolog phenotypic microarray are displayed and colored according to the difference in area between functional (WT) and non-functional (EXXQ) SMRs. Gray boxes indicate conditions where both functional and non-functional transporter were below the dead threshold (calculated from the area under the curve for non-functional SMR in PM15B, well 60) and analysis could not reliably be performed.

In our studies of EmrE, we chose four compounds with strong resistance or susceptibility hit scores for further phenotypic analysis: methyl viologen (MV), chelerythrine chloride (CC), harmane, and 18-crown-6 ether (18c6e) (34). Examining the hit scores for these 4 compounds across the panel of four SMR transporter shows some consistency between homologs. Methyl viologen and harmane are consistently strong resistance and susceptibility hits, respectively, for all four homologs (Fig. 2A, C; Table I). Chelerythrine chloride has a mixed phenotype with different behavior at low and high concentrations and a weaker phenotype for FTsmr than the other three homologs (Fig. 2B). 18-crown-6 ether (Fig. 2D) is a strong susceptibility hit for EmrE, FTsmr and SAsmr, but not for PAsmr. The full results (Fig. 1; Fig. S4) echo what we observe for these four compounds: all of the SMR homologs confer resistance to some compounds and susceptibility to others and while there is often consistency in the phenotype across different homologs for many substrates, there is also some variation in which compounds are substrates for each transporter. This variation in substrate specificity is expected given the sequence variation among the homologs, particularly in the TM1-3 region known to be important for determining the specificity profile of EmrE (20, 21, 35).

**Figure 2:**
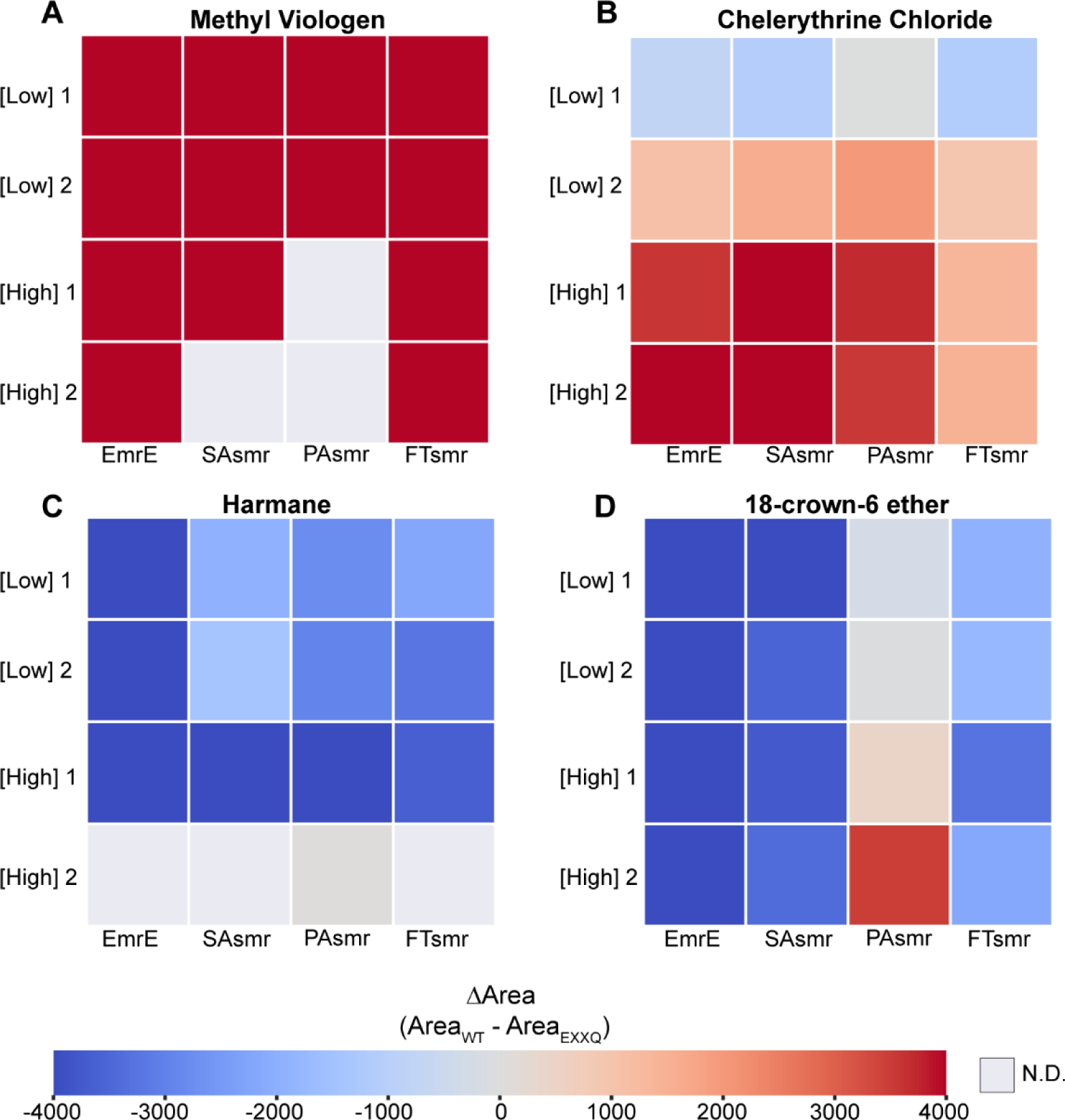
The functional promiscuity of SMR transporters extends beyond EmrE. Heat maps displaying the average ΔArea from two replicates of the Biolog assay for EmrE, SAsmr, PAsmr, and FTsmr. Data is shown for methyl viologen (A), chelerythrine chloride (B), harmane (C), and 18-crown-6 ether (D) with low and high concentration wells as indicated. Red indicates resistance and blue indicates susceptibility, with color intensity based on the strength of the phenotype. Gray boxes indicate conditions where both cell lines are below the dead threshold (calculated from the area under the curve for non-functional SMR in PM15B, well 60).

**Table I:** Summary of Biolog hit scores for specific compounds studied in this work.

| Compound | EmrE | SAsmr | PAsmr | FTsmr |
| --- | --- | --- | --- | --- |
| Methyl viologen | 8 | 6 | 4 | 8 |
| Chelerythrine Cl<sup>-</sup> | 6 | 5 | 6 | 4 |
| Harmane | -6 | -2 | -3 | -3 |
| 18-crown-6 ether | -5 | -5 | 1 | -2 |

Previous studies of EmrE substrate specificity suggested that substrate charge and hydrophobicity were key parameters affecting the affinity and transport rate EmrE substrates (20, 31, 50). These studies focused on known compound classes to which EmrE confers resistance (polyaromatic cations, quaternary ammonium compounds, etc.). Using our larger Biolog dataset, we compared the hydrophobicity (cLogP) and predicted charge for all the hits for EmrE (Fig. 3A), SAsmr (Fig. 3B), PAsmr (Fig. 3C), and FTsmr (Fig. 3D). Contrary to previous literature, hits from the Biolog microarray do not show any correlation between hydrophobicity and Biolog score for either susceptibility or resistance hits. However, charge does show some correlation with resistance vs susceptibility classification. Susceptibility hits for EmrE, SAsmr, and FTsmr were almost universally uncharged or only partially charged (pKa near 7) at neutral pH. In contrast, the majority of resistance hits were positively charged, as expected based on previous literature (31). However, a few resistance hits are likely to be negatively charged or neutral, and these will need to be confirmed as hits and the actual net charge of the substrates verified under the assay conditions. This trend is consistent across EmrE, SAsmr, and FTsmr confirming the previous characterization that SMR transporters confer resistance to hydrophobic cations, and that previous screens focusing on this compound class would have missed uncharged substrates to which SMR transporters confer susceptibility.

**Figure 3:**
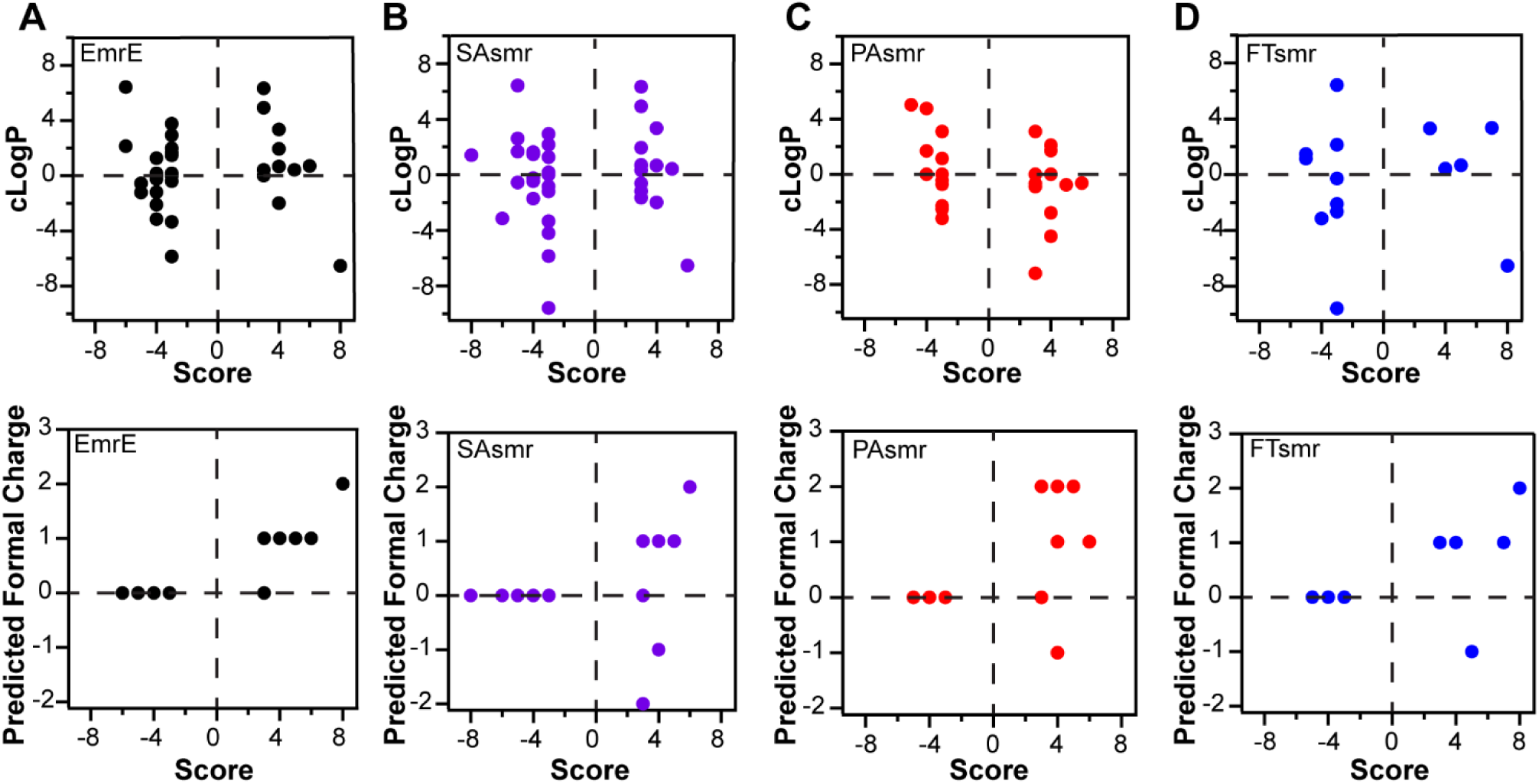
Substrate charge, not hydrophobicity, appears to be a determining factor in resistance versus susceptibility. The cLogP and formal charge of Biolog hits from EmrE (A), SAsmr (B), PAsmr (C), and FTsmr (D) reveal patterns that could differentiate resistance substrates from susceptibility substrates. The cLogP values and formal charges were taken from PubChem. There appears to be no influence of hydrophobicity on whether a substrate is a resistance or susceptibility hit of these SMR transporters. However, formal charge varies between the two groups of hits. Resistance hits (score ≥ 3) have a charge, whether it is positive or negative, but susceptibility hits (score ≤ -3) are neutral.

### Growth assays and dose-response curves confirm the metabolic results of the Biolog microarray

We monitored growth of MG1655 Δ*emrE E. coli* cells expressing each functional or non-functional SMR in the presence of methyl viologen (Fig. 4A-D), chelerythrine chloride (Fig. 4E- H), harmane (Fig. 5A-D), and 18-crown-6 ether (Fig. 5E-H) to test the robustness of the resistance or susceptibility phenotype for these four strong Biolog hits (Fig. 2) in an orthogonal assay. In the presence of methyl viologen and chelerythrine chloride (Fig. 4, S5A-F), MG1655 Δ*emrE E. coli* cells expressing functional SMR proteins (black) have increased growth compared to cells expressing their non-functional counterparts (red). Even though this resistance phenotype remains the same, the strength of the growth differential differs between the different SMR homologs. Since we are expressing all of the transporters in the same strain of *E. coli* and they express and insert in the membrane at comparable levels (Fig. S2), this differential reflects variation in the ability of the different SMR homologs to confer resistance to these compounds. Data for EmrE was previously published in (34) and is entirely consistent with the results of the Biolog assay. In general, these two assays measuring growth and metabolic activity are consistent for the other homologs as well. For example, FTsmr (Fig. 4H, S5F) has weaker resistance to chelerythrine chloride than the other homologs. However, there are a few exceptions – PAsmr has less resistance to methyl viologen and FTsmr has greater susceptibility to 18-crown-6-ether in the growth assay than expected based on the Biolog data. However, this could arise from differences in the exact compound concentration used in each of these assays.

**Figure 4:**
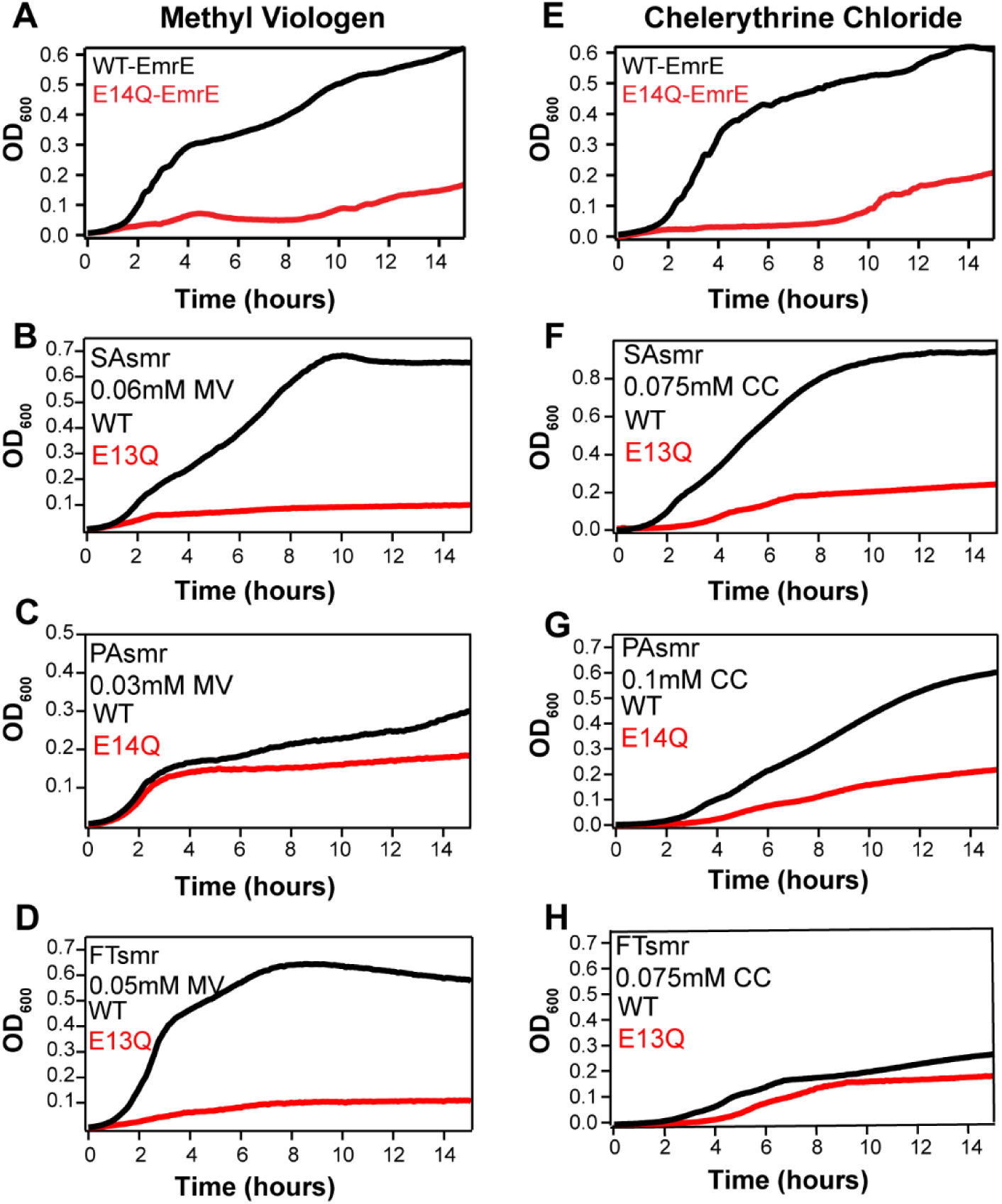
SMR transporters confer resistance to methyl viologen and chelerythrine chloride. Growth curves for functional (black) and non-functional (red) EmrE, SAsmr, PAsmr, and FTsmr expressed in MG1655 Δ*emrE E. coli* cells are displayed for methyl viologen (A-D) and chelerythrine chloride (E-H). The corresponding concentrations of each drug and the specific SMR are listed in each graph. While the resistance phenotype is present in all conditions, the strength of this resistance is different between each SMR homolog.

**Figure 5:**
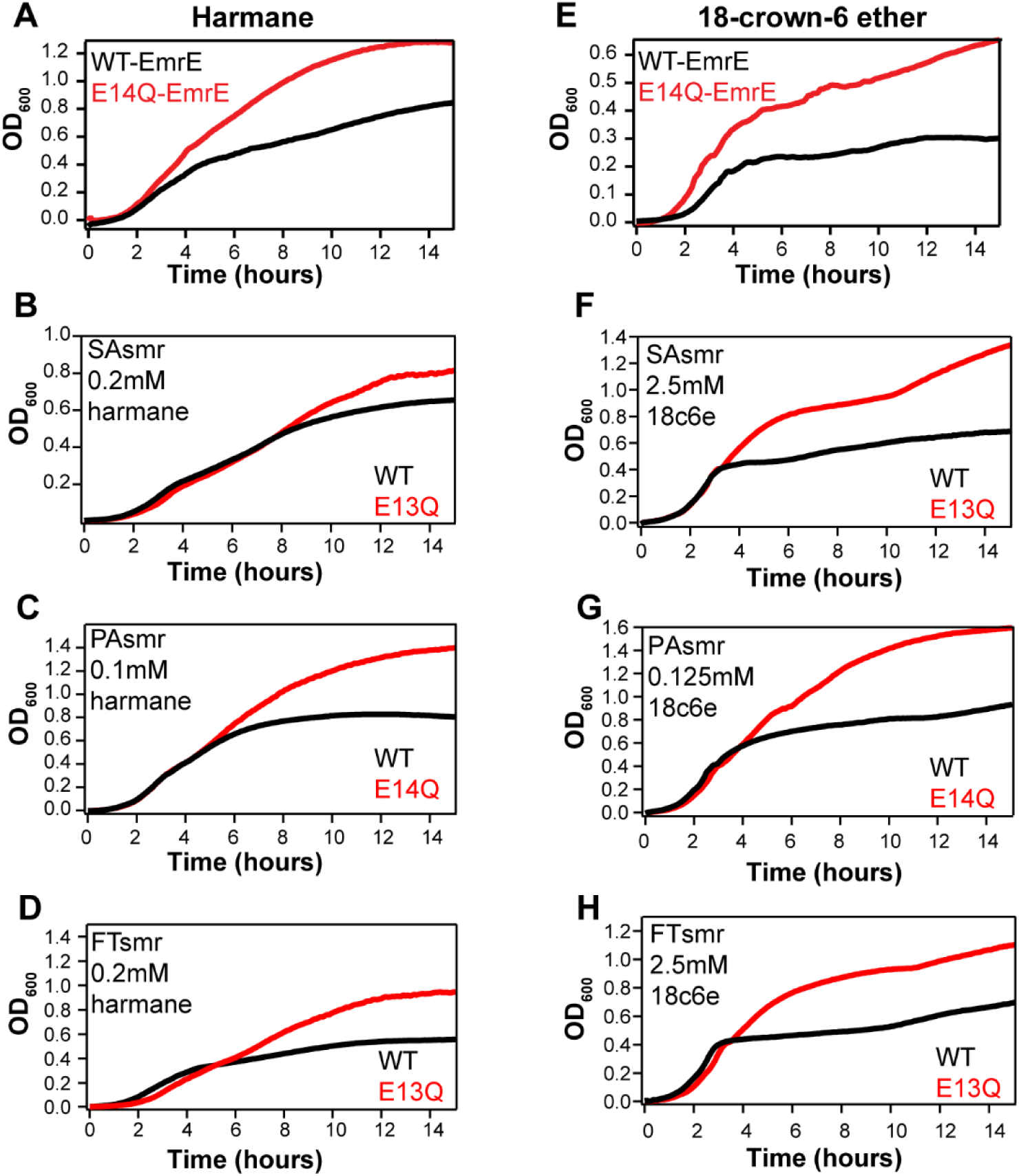
SMRs confer susceptibility to harmane and 18-crown-6 ether. Growth curves for functional (black) and non-functional (red) EmrE, SAsmr, PAsmr, and FTsmr expressed in MG1655 Δ*emrE E. coli* cells are displayed for harmane (A-D) and 18-crown-6 ether (E-H). The corresponding concentrations of each drug and the specific SMR are listed in each graph. Bacteriostatic phenotypes like those seen previously for EmrE occur a few hours after the assay is initiated, suggesting a metabolic link to the phenotype.

Dose-response curves of *E. coli* expressing EmrE, SAsmr, PAsmr, or FTsmr with each of these four compounds plus ethidium (known resistance substrate of EmrE and common MDR efflux pump substrate) and cetylpyridinium (Biolog resistance hit and antiseptic) further establish the functional behavior of these transporters (Table II, Figure S6A-D, S7A-D, S8A-D, S9A-D). IC_50_ values calculated from these curves show significant resistance to these compounds conferred by functional EmrE in the presence of ethidium, methyl viologen, chelerythrine chloride, and cetylpyridinium chloride (Fig. S6A-D). These results act as a benchmark for the ability of other SMRs to complement EmrE in *E. coli*. Functional SAsmr showed significant resistance to chelerythrine chloride (Fig. S7D), but minimal resistance to ethidium bromide, methyl viologen, and cetylpyridinium chloride (Fig. S7A, B, C), highlighting that the impact on growth and metabolism (Fig. 1, 2, 4, 5, S5) occurs in a narrow concentration window. This result is consistent with prior reports that SAsmr (*qacC*) conferred resistance to cetylpyridinium chloride and similar compounds in *S. aureus* (51). PAsmr is known to confer resistance to dyes and aminoglycosides in *P. aeruginosa* (38) along with the MexAB-OprM efflux system. Similarly, dose response curves and IC_50_ values reported here demonstrate that PAsmr confers significant resistance to ethidium bromide (Fig. S8A). Further, significant resistance to chelerythrine and cetylpyridinium is seen in *E. coli* expressing functional PAsmr (Fig. S8C, D), but little resistance to methyl viologen (Fig. S8B). There is little information available on the function of FTsmr *in vivo*, but we see significant, consistent resistance to methyl viologen in *E. coli* expressing functional FTsmr (Fig. S9B). IC_50_ values increase by 2- fold for *E. coli* expressing WT-FTsmr for ethidium, chelerythrine, and cetylpyridinium. While this 2-fold increase is not significant according to our thresholds, it demonstrates that even FTsmr can complement EmrE in *E. coli*.

**Table II:**
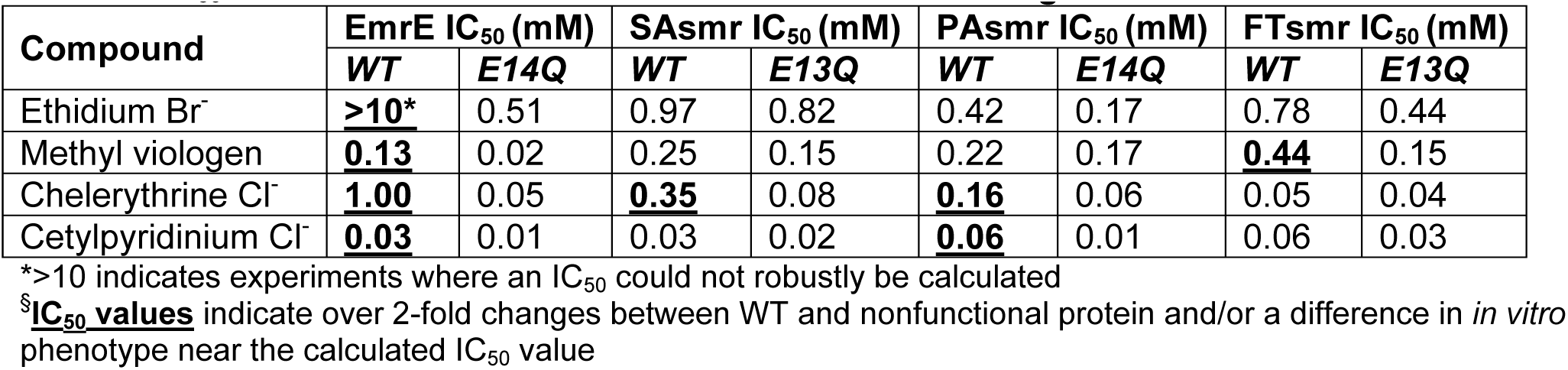
IC_50_ values of resistance hits of EmrE and its homologs

| Compound | EmrE IC<sub>50</sub> (mM) |  | SAsmr IC<sub>50</sub> (mM) |  | PAsmr IC<sub>50</sub> (mM) |  | FTsmr IC<sub>50</sub> (mM) |  |
| --- | --- | --- | --- | --- | --- | --- | --- | --- |
|  | WT | E14Q | WT | E13Q | WT | E14Q | WT | E13Q |
| Ethidium Br<sup>-</sup> | **>10\*** | 0.51 | 0.97 | 0.82 | 0.42 | 0.17 | 0.78 | 0.44 |
| Methyl viologen | **0.13** | 0.02 | 0.25 | 0.15 | 0.22 | 0.17 | **0.44** | 0.15 |
| Chelerythrine Cl<sup>-</sup> | **1.00** | 0.05 | **0.35** | 0.08 | **0.16** | 0.06 | 0.05 | 0.04 |
| Cetylpyridinium Cl<sup>-</sup> | **0.03** | 0.01 | 0.03 | 0.02 | **0.06** | 0.01 | 0.06 | 0.03 |

\*>10 indicates experiments where an IC<sub>50</sub> could not robustly be calculated
<sup>§</sup>IC<sub>50</sub> values indicate over 2-fold changes between WT and nonfunctional protein and/or a difference in *in vitro* phenotype near the calculated IC<sub>50</sub> value

The more novel result, that SMR homologs can confer susceptibility to some substrates rather than resistance, was also further assessed by measuring dose-response curves for each SMR with harmane or 18-crown-6 ether (Table III, Figure S6E,G; S7E,G; S8E,G; S9E,G). Dose response curves for EmrE with harmane (Fig. S6E) or 18-crown-6 ether demonstrates a significant decrease in IC_50_ values when *E. coli* express functional EmrE (Table III, Fig. S6G). Although harmane had a stronger, more consistent susceptibility phenotype in the Biolog assay (Fig. 2), only EmrE show a consistent reduction in harmane IC_50_ values with functional transport. In contrast, 18-crown-6 ether was a weaker, less-consistent hit in the Biolog assay but there is a significant decrease in IC_50_ values for this substrate with functional EmrE (Fig. S6G), SAsmr (Fig. S7G) and PAsmr (Fig. S8G). While there is less consistency between assay formats for the susceptibility hits, these results still demonstrate that EmrE and its homologs from other SMRs can confer susceptibility to some substrates *in vivo*.

**Table III:**
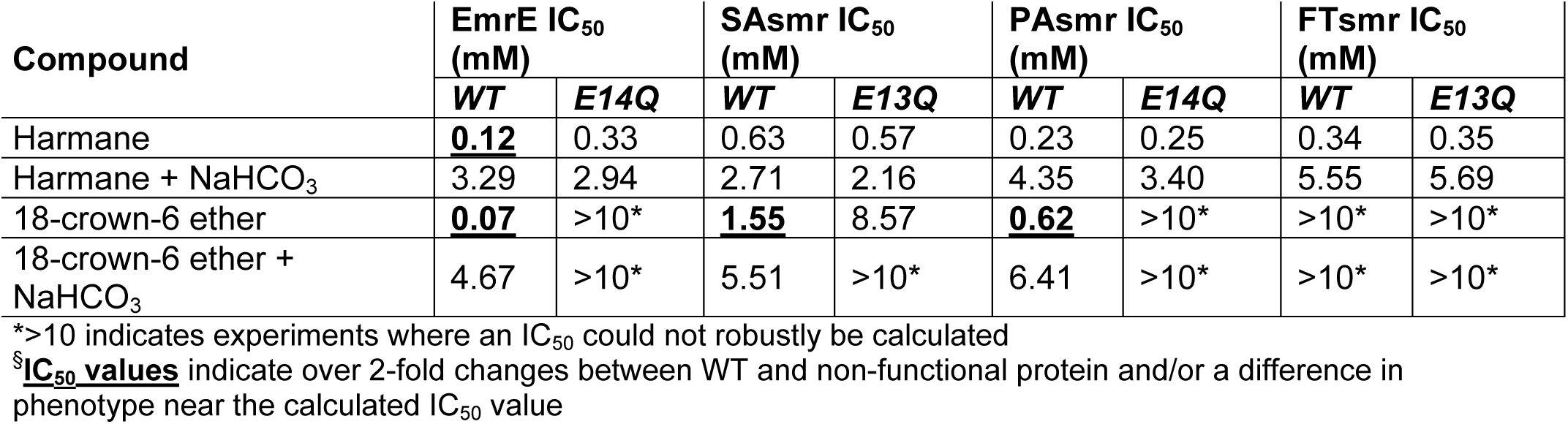
IC_50_ values of susceptibility hits of EmrE and its homologs

| Compound | EmrE IC<sub>50</sub><br>(mM) |  | SAsmr IC<sub>50</sub><br>(mM) |  | PAsmr IC<sub>50</sub><br>(mM) |  | FTsmr IC<sub>50</sub><br>(mM) |  |
| --- | --- | --- | --- | --- | --- | --- | --- | --- |
|  | *WT* | *E14Q* | *WT* | *E13Q* | *WT* | *E14Q* | *WT* | *E13Q* |
| Harmane | **0.12** | 0.33 | 0.63 | 0.57 | 0.23 | 0.25 | 0.34 | 0.35 |
| Harmane + NaHCO<sub>3</sub> | 3.29 | 2.94 | 2.71 | 2.16 | 4.35 | 3.40 | 5.55 | 5.69 |
| 18-crown-6 ether | **0.07** | >10* | **1.55** | 8.57 | **0.62** | >10* | >10* | >10* |
| 18-crown-6 ether + NaHCO<sub>3</sub> | 4.67 | >10* | 5.51 | >10* | 6.41 | >10* | >10* | >10* |

\*>10 indicates experiments where an IC<sub>50</sub> could not robustly be calculated709 <sup>§</sup>**IC<sub>50</sub> values** indicate over 2-fold changes between WT and non-functional protein and/or a difference in phenotype near the calculated IC<sub>50</sub> value

These results show some variation in specific substrate profile for each SMR homolog, as expected based on sequence variation in regions of the transporter known to be important for substrate binding and specificity. Although the results of the growth and metabolic assays characterizing the ability of SMR transporters to confer susceptibility or resistance to different compound in *E. coli* are not perfectly consistent, these orthogonal assays confirm that the functional promiscuity previously described for EmrE (34) does extend to at least three other SMR homologs across the same subfamily.

### Bicarbonate eradicates the phenotype of harmane and 18-crown-6-ether for all tested SMRs

We previously demonstrated that EmrE confers susceptibility to harmane because harmane binds to the transporter and triggers uncoupled proton flux (34). In other words, EmrE acts as a harmane-gated proton uniporter, but it functions as a proton-coupled antiporter of ethidium and other resistance substrates. Uncontrolled proton leak through EmrE dissipates the ΔpH component of the proton motive force, leading to the specific growth and metabolic defects detected in the *in vivo* assays. This is consistent with the time-dependent bacteriostatic effects observed in the growth curves for harmane-induced susceptibility with functional SMR transporter, while methyl viologen has a more immediate and complete suppression of bacterial growth unless functional transporter is present to confer resistance. In addition, bacteria compensate for ΔpH dissipation by enhancing Δψ (membrane potential) to attempt to maintain a stable proton motive force (52), which may explain the more variable phenotype observed in the different assays for the susceptibility hits.

To test for ΔpH dissipation *in vivo*, we repeated the dose-response curves for the susceptibility hits (harmane and 18-crown-6-ether) in the presence of 25 mM sodium bicarbonate. Sodium bicarbonate can diffuse directly through the membrane and dissipates ΔpH independently of any membrane protein or transporter. This will suppress the susceptibility phenotype if it is due to proton leak since there will no longer be any ΔpH to dissipate. This is exactly what was previously observed for EmrE (34), and is replicated here (Table III, Fig. S6F, H). None of the other transporters showed a susceptibility phenotype in the dose response curves with harmane and bicarbonate, and addition of bicarbonate increases the IC_50_ value equally for both functional and non-functional transporter. 18-crown-6 ether showed a more consistent susceptibility phenotype in *E. coli* expressing the various SMR homologs. In the presence of 25mM sodium bicarbonate, the IC_50_ values for 18-crown-6-ether increase by 5-fold or more (Table III, Fig. S6H, S7H, S8H, S9H), showing a strong suppression of the impact of SMR activity. These results are consistent with substrate-gated proton leak and ΔpH dissipation as mechanism of harmane- and 18-crown-6-ether-induced susceptibility.

### Harmane acts as an antibiotic adjuvant in the presence of SMRs

With the rising rates of antibiotic resistance, there is increasing interest in finding chemical adjuvants to include in existing antibiotic treatments. Since harmane dissipates ΔpH, it should synergize with compounds that dissipate Δψ, resulting in a more significant impact on the total PMF. Bacteria maintain the total PMF, compensating for reduced ΔpH by increasing Δψ. Thus, antibiotics with Δψ-dependent uptake or activity should also have enhanced efficacy in the presence of harmane due to the natural bacterial response upon EmrE-mediated dissipation of ΔpH. There is some evidence that cellular uptake of aminoglycosides requires Δψ, while tetracycline and other similar antibiotics require the ΔpH component of the proton motive force to enter the cell (53). We previously performed checkerboard assays of *E. coli* expressing WT-EmrE treated with harmane and either kanamycin or tetracycline (34). Harmane was shown to synergize with kanamycin, but not with tetracycline, consistent with our hypothesis for ΔpH dissipation by harmane. To complete our study of EmrE homologs from *S. aureus*, *P. aeruginosa*, and *F. tularensis*, we performed the same checkerboard assays with *E. coli* expressing WT and non-functional SMR (Fig. 6, 7). Functional WT-SAsmr, PAsmr and FTsmr displayed synergy between harmane and kanamycin (Fig. 6), similar to EmrE. In contrast, but again consistent with EmrE (34) there is no observable interaction between harmane and tetracycline (Fig. 7), for any of the SMR homologs tested. These results demonstrate synergy between harmane and kanamycin in a manner that requires active SMR transporter and are consistent with the proposed mechanism of ΔpH dissipation.

**Figure 6:**
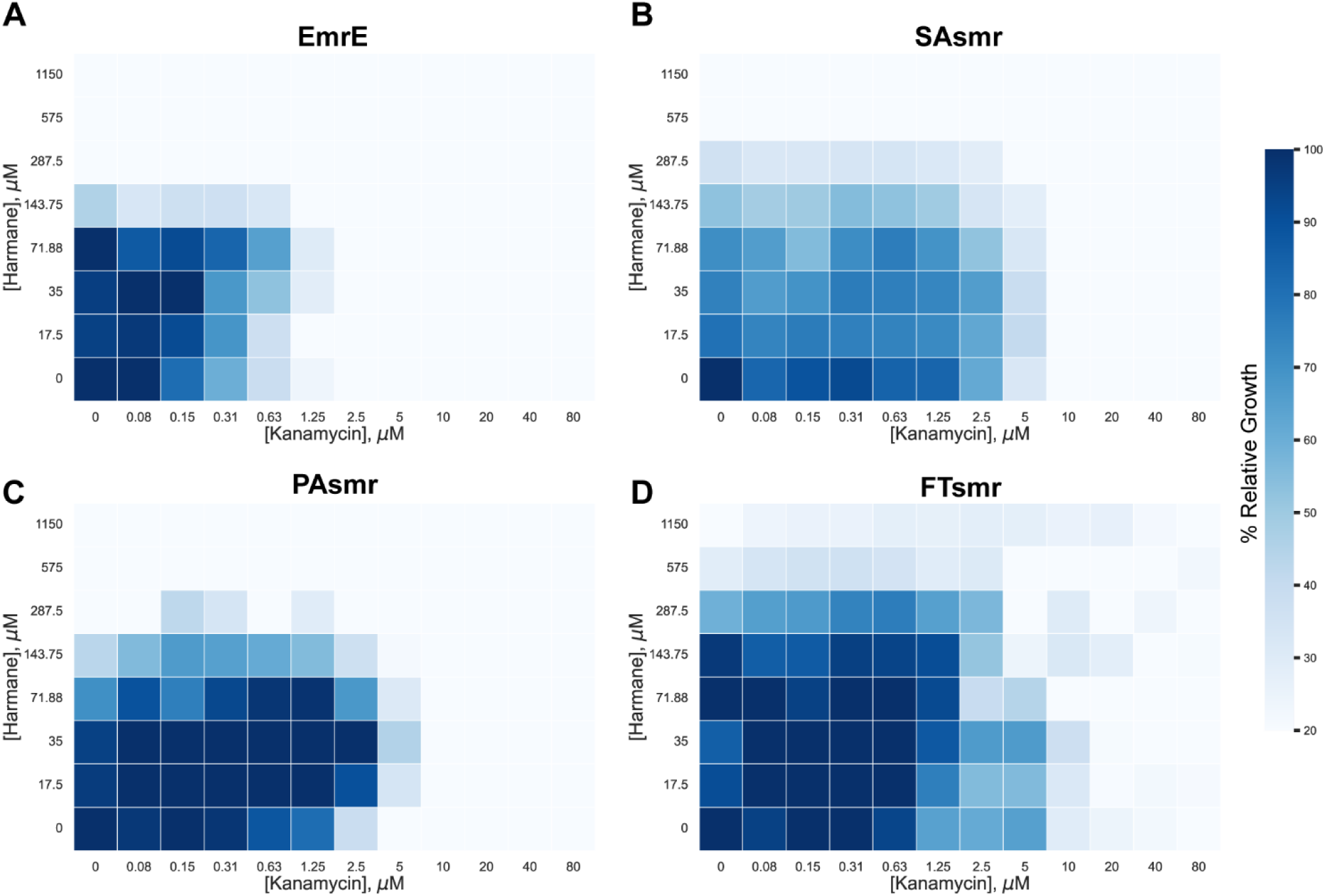
Harmane acts as an adjuvant with kanamycin. Checkerboard assays of MG1655 Δ*emrE E. coli* cells expressing WT-EmrE (A), SAsmr (B), PAsmr (C), or FTsmr (D) show that harmane acts as an adjuvant for kanamycin and this synergy requires functional transporter (Fig. S10). Each cell represents the average relative growth across three technical replicates. The data is colored by % relative growth compared to no drug (lower left well) as measured by OD_600_, with relative growth at or below 0.2 (80% inhibition) is shown in white.

**Figure 7:**
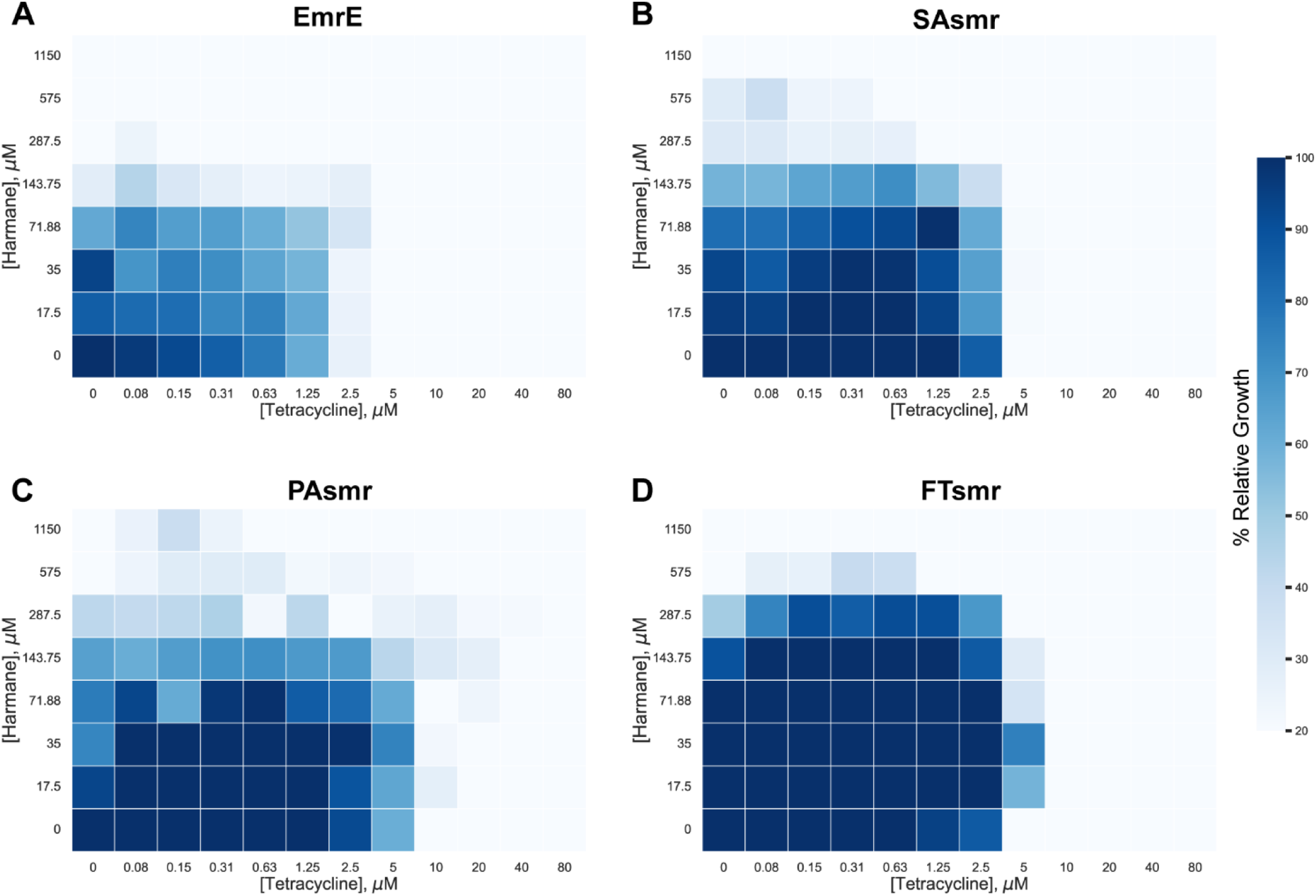
Harmane interacts differently with tetracycline. Checkerboard assays of MG1655 Δ*emrE E. coli* cells expressing WT-EmrE (A), SAsmr (B), PAsmr (C), or FTsmr (D) show that harmane interacts differently when used as a co-treatment with tetracycline. Each cell represents the average relative growth across three technical replicates. The data is colored by % relative growth compared to no drug (lower left well) as measured by OD_600_, with relative growth at or below 0.2 (80% inhibition) is shown in white.

## DISCUSSION

The increasing rates of antibiotic resistance across bacterial species from *E. coli* (54), *Staphylococcus* (44, 51), *P. aeruginosa* (38, 46), and *F. tularensis* (40), among others, necessitate the formulation of novel antibiotic therapeutic strategies. Previous work on EmrE demonstrates that this SMR transporter can be triggered by harmane to rundown the ΔpH component of the proton motive force in *E. coli* to the extent that it has a negative impact on bacterial growth and metabolism (34). In this work, we demonstrate that these results extend to EmrE homologs across the promiscuous EmrE-like subfamily of SMR transporters, including those from pathogenic strains of bacteria: SAsmr, PAsmr, and FTsmr. Biolog phenotypic microarrays (Fig. 1, 2), growth curves (Fig. 4, 5, S5), and dose-response curves (Table II and III; Fig. S6, S7, S8, S9) demonstrate consistently that SMR and its homologs can confer either resistance or susceptibility *in vivo*, depending on the identity of the small molecule substrate. Replication of these *in vivo* experiments in the endogenous strains for these SMRs and broader substrate screening to better define the molecular properties that distinguish resistance and susceptibility substrates and identify stronger susceptibility hits are the logical next steps to assess whether the SMR transporter family provide a viable target for development of antibiotic adjuvants.

Inhibition of SMRs has been studied previously as a new strategy to combat resistant bacteria (55, 56), but SMR family members are rarely the dominant efflux pump contributing to antibiotic resistance in bacteria. Thus, simple inhibition is not likely to have a major clinical impact. We do not propose to inhibit SMR-mediated efflux, but rather shift these transporters from proton-coupled-substrate-antiport to substrate-triggered-proton-leak, effectively using small molecules to drive an entirely different transport function. This alternative function dissipates ΔpH, impacting bioenergetics. It is unlikely that ΔpH dissipation will ever provide the bactericidal efficacy desired for an antibiotic, but it has potential as antibiotic adjuvant for two reasons. First, as demonstrated through the synergy with kanamycin shown here (Fig. 6), it can synergize with existing antibiotics whose activity is influenced by the membrane potential. Second, the major antibiotic efflux pumps in bacteria are proton-coupled antiporters, and dissipation of ΔpH to the extent that it impacts bacterial growth and metabolism, as demonstrated here, will reduce the energy source for antibiotic resistance through these other proton-coupled pumps. Usage of antibiotic adjuvants is increasing as standard antibiotics are losing their efficacy. To date, the only FDA-approved adjuvants are beta-lactamase inhibitors, but other classes are in development (57). Some of these classes target efflux pumps for inhibition or to act as routes of concentrative uptake of antibiotics in bacteria. This work suggests that the SMR transporter family may represent an alternative target for development of antibiotic adjuvants with a novel mechanism of action.

## MATERIALS AND METHODS

### Bacterial Strains and Plasmids

MG1655 Δ*emrE E. coli* cells were transformed with either WT or non-functional (E13/14Q) versions of EmrE (AAN90039.1), SAsmr (QacC) (WP_031824198), PAsmr (Q9HUH5), and FTsmr (ABK89687.1) cloned into the pWB plasmid, a low copy, leaky-expression vector. These plasmids were used in all assays described in this paper.

### Sequence alignment

A sequence alignment was generated for the accession numbers of EmrE, SAsmr, PAsmr, and FTsmr using Clustal Omega (58–60). The phylogenetic tree of all SMR transporters identified in the Pfam subfamily (13) was calculated using Clustal Omega and visualized using the Interactive Tree of Life (iTOL) software (1, 61).

### Biolog Phenotypic Microarrays

MG1655 *ΔemrE E. coli* cells containing either WT- or E13/14Q-SMR (non-functional) constructs from the bacterial strains used in this paper were grown on LB-Carb media overnight at 37°C. The phenotype microarray tests followed the established protocols of standard PM procedures for *E. coli* and other gram-negative bacteria (12). PM01-20 plates were used to screen both WT- and non-functional SMR expressing cells. Overnight plates were resuspended in IF-0a inoculating fluid (Biolog) to an optical density of 0.37. The cells were diluted by a factor of 6 into IF-0a media plus Redox Dye A and 20mM glucose was added for PM3-8 plates. Cells were diluted to a 1:200 dilution in IF-10a media (Biolog) with Redox Dye A for PM9-20 plates. PM plates were inoculated with 100µL of cell suspensions per well. The microplates were incubated at 37°C and read using the OmniLog instrument every 15min for 24 h. The area under the resulting metabolic curves was determined for cells expressing WT-SMR or E13Q/E14Q- SMR. The difference was calculated as:

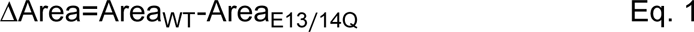

Resulting in positive values for greater respiration by cell expressing WT-SMR and negative values for greater respiration by cells expressing non-functional SMR. The 10% trimmed mean was then calculated for each data set (ΔArea replicate 1, ΔArea replicate 2) separately for each transporter as variation between replicates can arise due to minor deviations between plate sets or in the exact concentration of dye or OD of cells upon dilution on different days. The standard deviation was then calculated among known non-hits (selecting at least 50 wells out of the 960 total wells in a single data set) to determine the cut-off values for actual hits. Individual wells were assessed as hits if the calculated ΔArea value (equation 1) was more than two standard deviations from the 10% trimmed mean. For each hit, a value of +1 was assigned for resistance hits (positive ΔArea) and a value of -1 was assigned for susceptibility hits (negative ΔArea). These values were then summed across all eight wells for a single compound (4 wells of the same compound per plate set * 2 replicates, with a max score of ±8. Final resistance or susceptibility hits were assigned if the total score was ≥ +3 (resistance) or ≤ -3 (susceptibility). This definition was chosen since small total hit scores of ±1 or ±2 could arise by chance using the ± 2*SD cutoff to score individual wells. Values of ±3 recognize consistent hits across multiple replicates and/or different concentrations of the same compound. Our cutoff is not set higher since the 4 wells of each compound on a single plate set include different concentrations and some concentrations may not be sufficient to elicit a phenotype.

### Microplate Growth Curves

MG1655 Δ*emrE E. coli* expressing either WT or non-functional forms of EmrE, SAsmr, PAsmr, and FTsmr were grown in Mueller Hinton Broth (100 µg/mL carbenicillin) from a single colony to an OD of 0.2 at 37 °C. The cells were then diluted to a final OD of 0.01 in microplates (Corning, REF: 351172) containing select concentration values of Eth^+^, MV^2+^, chelerythrine chloride, cetylpyridinium chloride, harmane, and 18-crown-6 ether as determined by the SMR being tested. The plates were incubated in a microplate reader (BMG-Labtech) at 37 °C. OD_600_ (or OD_700_ for Eth^+^) were measured every 5 minutes for 20 hours. Experiments were performed in technical and biological triplicate and data was analyzed using Excel and Igor Pro.

### Dose-response curves

MG1655 Δ*emrE E. coli* expressing either WT or non-functional forms of EmrE, SAsmr, PAsmr, and FTsmr were grown in Mueller Hinton Broth (100 µg/mL carbenicillin) from a single colony to an OD of 0.2 at 37 °C. Cells were diluted to a final OD of 0.01 in 96-well microplates with three biological replicates. Serial dilutions of the drugs from the microplate assays (Eth^+^, MV^2+^, chelerythrine chloride, cetylpyridinium chloride, harmane, and 18-crown-6 ether) in 1x MHB were added to the wells with two blank replicates (no cells) per drug concentration. Plates were incubated at 37 °C and shaken at 300 rpm in a microplate incubator. OD_600_ (or OD_700_ for Eth^+^) was measured after 18 hours by a BMG-Labtech plate reader. Replicates were averaged, and blank measurements were subtracted from the respective OD_600_. Relative growth was calculated by dividing the averaged OD_600_ of each concentration by the averaged OD_600_ with no drug.

### Bicarbonate synergy/antagonism assays

MG1655 Δ*emrE E. coli* containing plasmids of interest were grown up in Mueller Hinton Broth (100 µg/mL carbenicillin) from a single colony to an OD of 0.2 at 37 °C. Cells were diluted to a final OD of 0.01 in 96-well microplates with three biological replicates. Varying concentrations of harmane and 18-crown-6 ether in 1x MHB were added to wells with 25mM bicarbonate (pH 7.4) and two blank replicates (no cells) per concentration. Plates were incubated at 37 °C and shaken at 300 rpm in a microplate incubator. OD_600_ was measured after 18 hours by a BMG- Labtech plate reader. Replicates were averaged, and blank measurements were subtracted from the respective averaged OD_600_ with no drug.

### Checkerboard assays

MG1655 *ΔemrE E. coli* cells expressing either WT- or nonfunctional E13Q constructs of EmrE, SAsmr, PAsmr, and FTsmr were grown in Mueller-Hinton broth (Sigma, 100µg/mL carbenicillin, pH 7.0) from a single colony to an OD of 0.2 at 37 °C. Kanamycin or tetracycline was serially diluted across a 96-well microplate in MHB with concentrations ranging from 0- 80μM or 0-16μM respectively. Either harmane or 18-crown-6 ether were serially diluted down a separate plate using MHB with concentrations ranging from 0-1150μM. The solution from this plate was then transferred to the corresponding wells in the first plate to reach the final concentrations listed above. Cells were then diluted to a final OD of 0.01 in the microplate. A column with no kanamycin or tetracycline and a row with no harmane was used to determine the MIC values for each compound. Inoculated plates were sealed and incubated with shaking for 18h at 37°C. OD_600_ endpoints were taken using a microplate reader (BMG-Labtech). Checkerboard synergy testing was performed in triplicate and analyzed for MIC and FIC values in Excel.

The fractional inhibitory concentration (FIC) index was calculated for each well with no turbidity along the interface using the MIC values for the different compounds individually and in tandem. The MIC value was defined as the minimum concentration required to inhibit all cell growth to 20% of the background growth, as detailed in (62). FIC values were determined using the following equations:

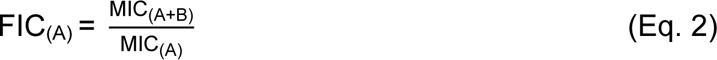

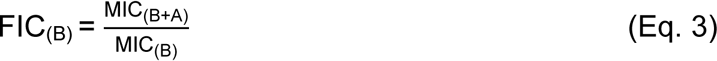

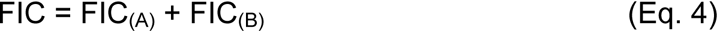

where A and B represent the different compounds in the assay. The FIC index for each drug combination was calculated for a well showing <20% growth and used to determine synergism (FIC < 0.5), indifference (0.5 ≤ FIC < 1), or antagonism (FIC ≥ 1).

## SUPPLEMENTARY MATERIAL

Supplemental materials are available online only. Supplemental file 1

Fig. S1-S11, and reference to data repository.

Data repository (raw *in vivo* data and Python scripts): <u>MendeleyData</u> (when published, update to http://doi.org/10.17632/vs6wfsfdtm.1)

## ACKNOWLEDGEMENTS

The authors would like to thank Jason Peters for his intellectual contribution to this manuscript. The authors wish to thank Grant A. Hussey for aiding in the initial plasmid design of the pWB vector and Kylie M. Hibbs for preliminary growth curve work. Thanks also to Trey Sato for his assistance in using the OmniLog Instrument for Biolog Assays with funding from the Great Lakes Bioenergy Research Center under U.S. Department of Energy, Office of Science, Office of Biological and Environmental Research Award Number DE-SC0018409. This work was supported by the USDA National Institute of Food and Agriculture Hatch WIS01985. Research reported in this publication was supported by the institute for General Medical Sciences of the National Institutes of Health under award number R01GM095839 and R35GM141748. The content is solely the responsibility of the authors and does not necessarily represent the official views of the NIH, USDA, or NIFA.

## AUTHOR CONTRIBUTIONS

PJS – Biolog assays, Manuscript Drafting/Editing, Figures, Conception/Design, Interpretation, Supervision. CJP – Growth curves, IC_50_ values, Checkerboard Assays, Assay Design, Interpretation, Figures, Manuscript Editing. AW – Cloning, Expression validation, Manuscript Editing, Supervision. WFB – Cloning, Experimental Design, Preliminary Data. SPD and ENP – Experimental replicates for growth curves, IC_50_ values, and checkerboard assays, Manuscript Editing. KAHW – Funding Acquisition, Supervision, Conception/Design, Manuscript Drafting/Editing, Interpretation.

## Supplemental Materials

Additional datasets and scripts for heat maps can be found at: https://doi.org/10.17632/vs6wfsfdtm.1

Until this paper is published, please use the following link: https://data.mendeley.com/datasets/vs6wfsfdtm/draft?a=ee4d52cf-c338-4a7b-8b66-34cc50bd1fd1

**Figure S1:**
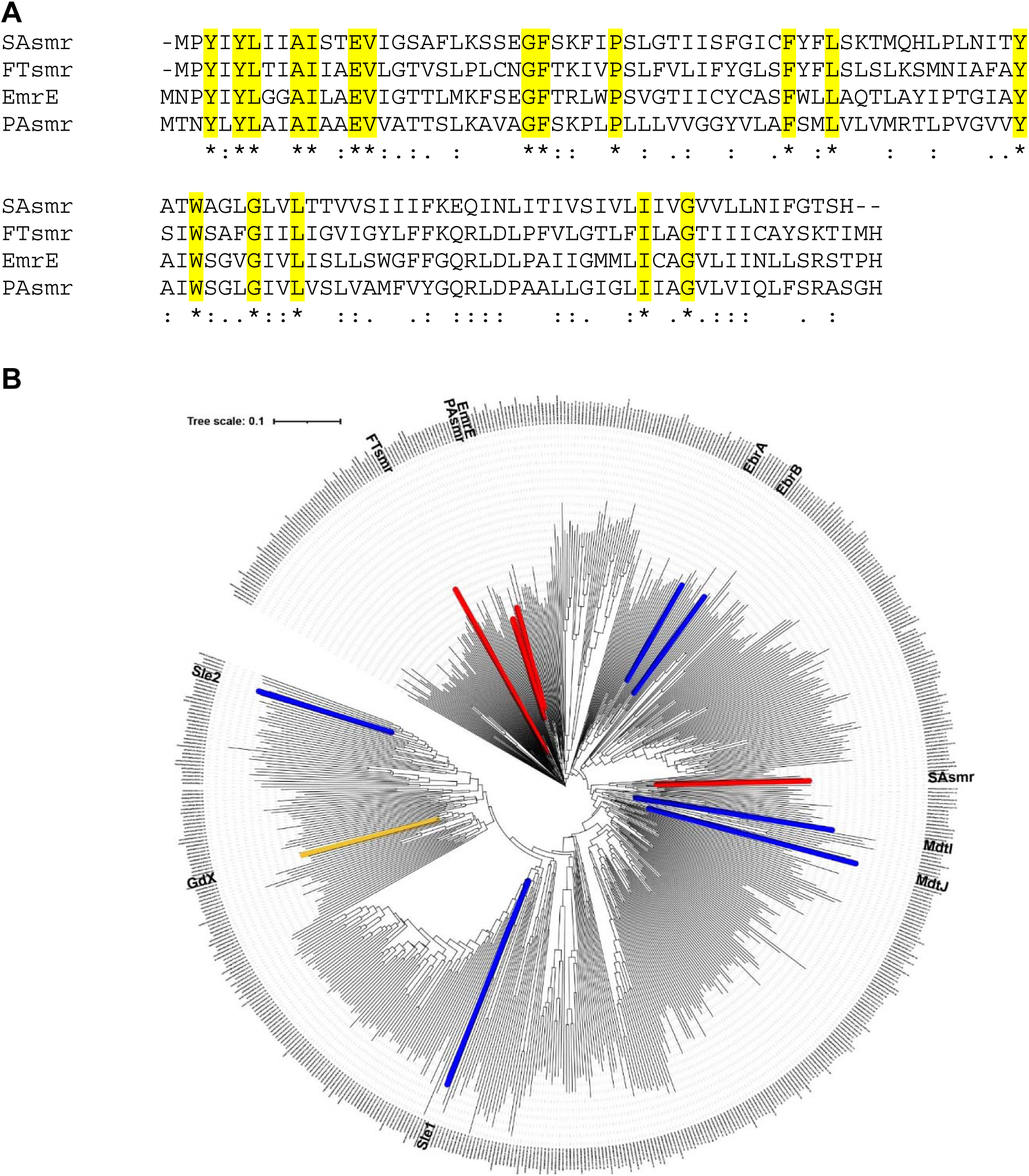
Sequence identity of EmrE and SAsmr, PAsmr, and FTsmr. A) Alignment of EmrE, SAsmr, PAsmr, and FTsmr reveals 40-45% sequence identity for SAsmr, PAsmr, and FTsmr compared to EmrE. (B) However, these transporters are relatively diverse within the promiscuous EmrE-like multidrug subfamily of SMR transporters. Promiscuous drug efflux homodimer SMR transporters tested in this manuscript are highlighted in red. GdX, a specific guanidinium transporter is highlighted in yellow, showing which region of the tree corresponds to the more specific Gdx-like metabolite exporters. Paired multidrug efflux SMR transporters are highlighted in blue and are found throughout both the EmrE-like and Gdx-like subfamilies. This tree was constructed using the multiple sequence alignment from Kermani et al., 2018 (DOI: 10.1073/pnas.1719187115).

**Figure S2:**
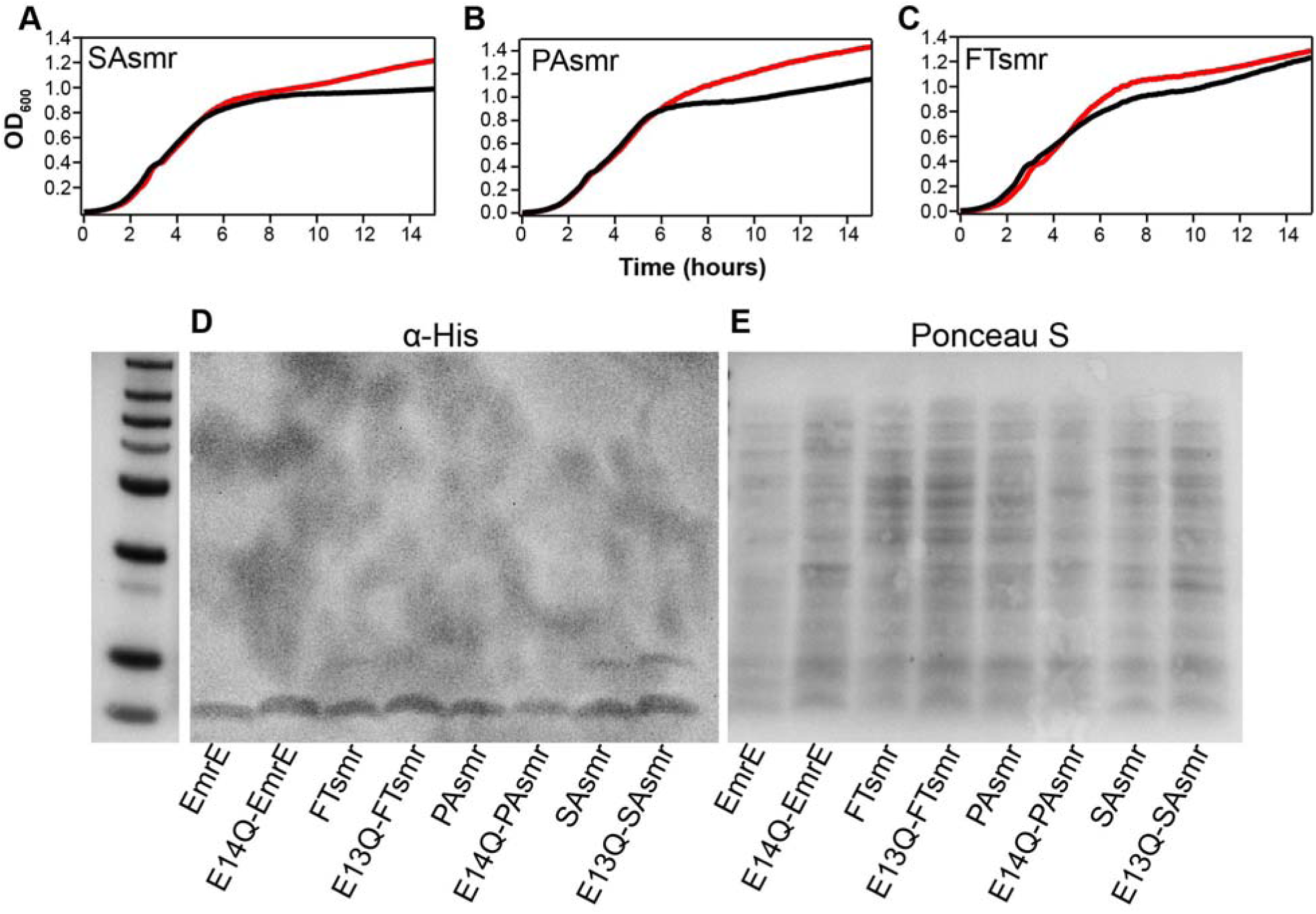
Growth and expression of functional and non-functional SMR homologs. A-C) Leaky expression of the SMR homologs from a low copy number plasmid in MG1655 Δ*emrE E. coli* shows minimal impact on bacterial growth for expression of functional (black) versus non-functional (E13Q- or E14Q-) mutants (red), and similar growth with all of the different homologs in the absence of any small molecule substrates. (D, E) Western blot to the His-tag on N-terminal His-tagged versions of each transporter reveals equivalent expression across all SMR homologs and their non-functional mutants (D). Ponceau S (E) was used as a loading control.

**Figure S3:**
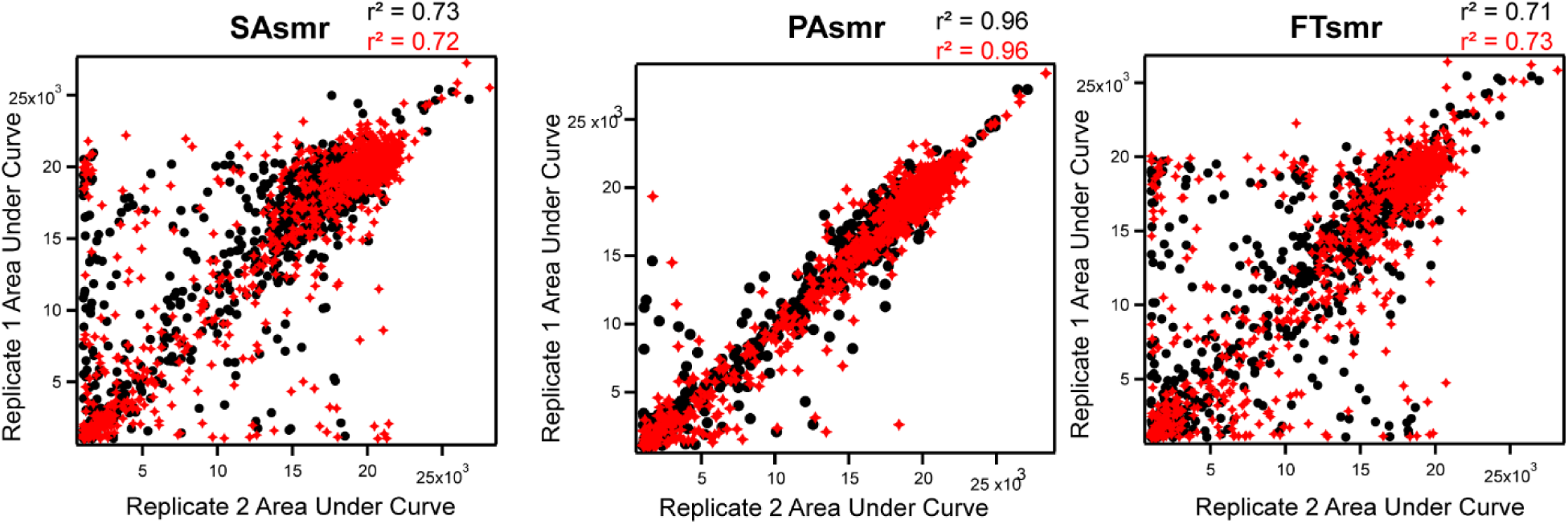
Correlation between replicates of Biolog microarrays. The correlation of the area under the curve between individual replicates of SAsmr, PAsmr, and FTsmr (WT, black; non-functional, red) are displayed above. Correlation of EmrE replicates has been previously published (Spreacker et al., 2022, *BioRxiv*). The high correlation of these data (r^2^ values) make analysis of the Biolog phenotypic microarray more robust.

**Figure S4:**
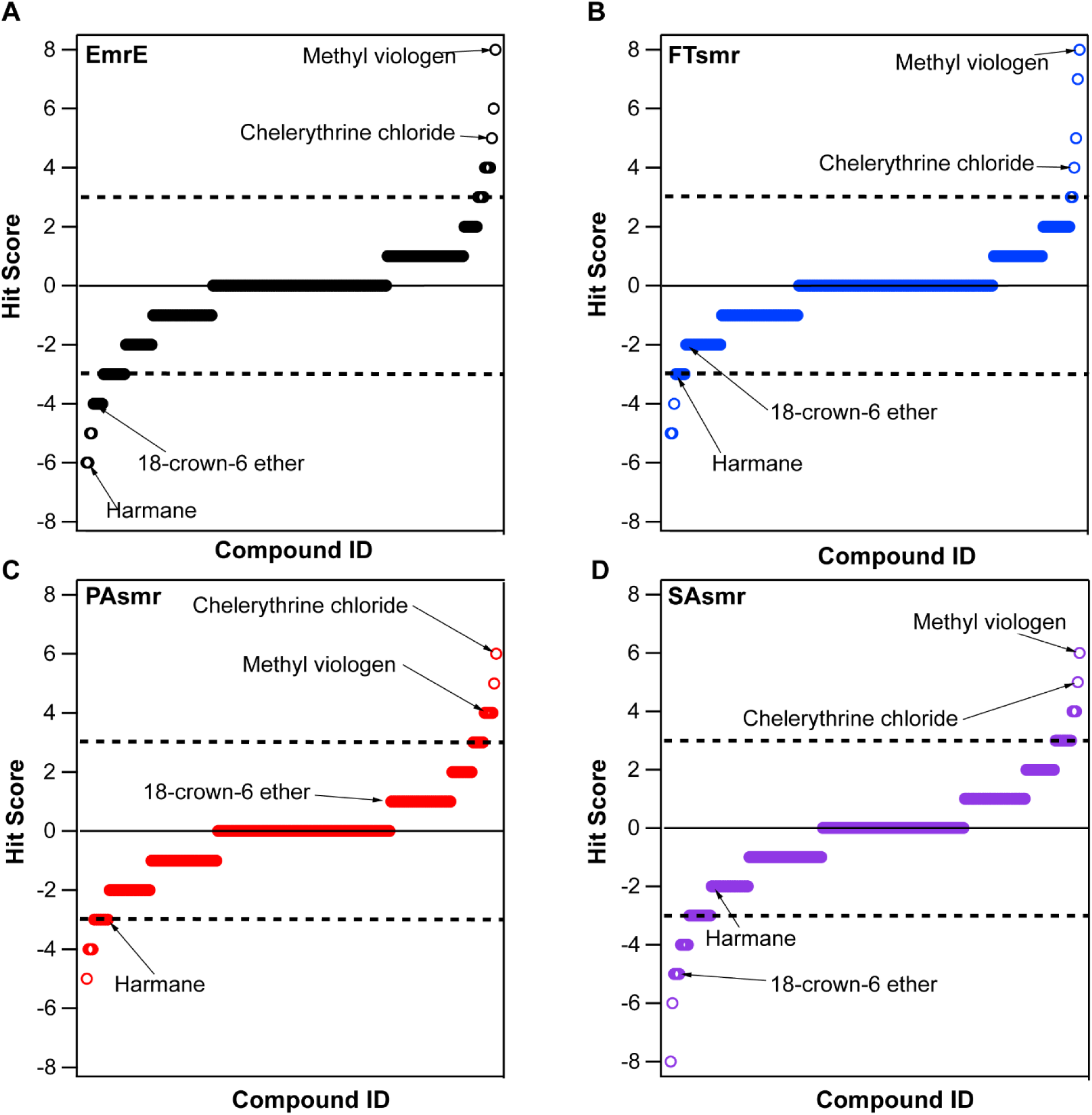
Biolog Data by SMR transporter sorted by hit score. Hit scores range from +8 (all wells significant resistance phenotype) to -8 (all wells significant susceptibility phenotype). Data is show separately for A) EmrE B) FTsmr, C) PAsmr, and D) SAsmr (purple), The four compounds used in further experimentation are denoted with an arrow.

**Figure S5:**
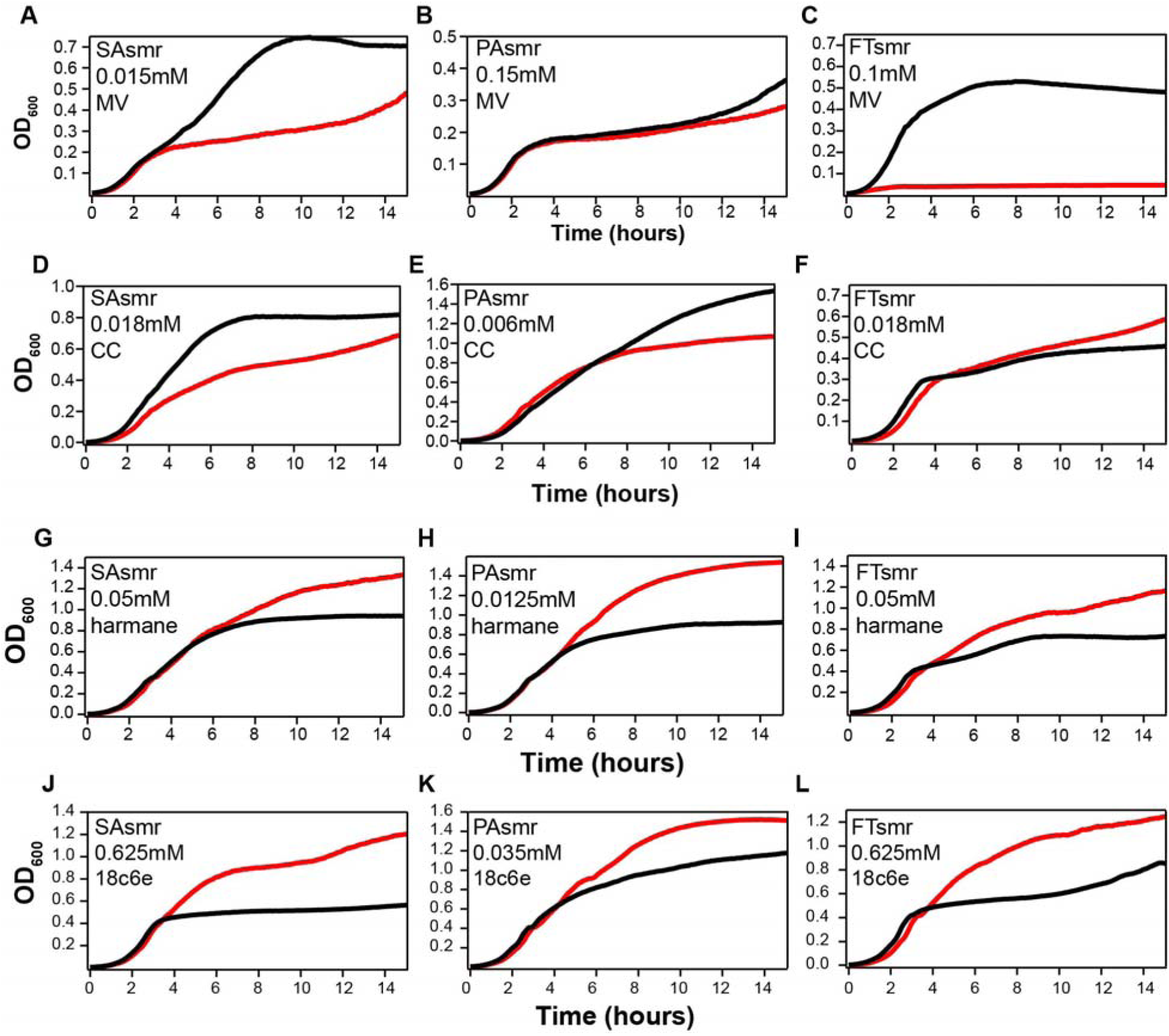
Additional growth curves for methyl viologen, chelerythrine chloride, harmane, and 18-crown-6 ether. Additional growth curves for MG1655 Δ*emrE E. coli* expressing functional (black) or non-functional (red) SMR homologs at various substrate concentrations for methyl viologen (A-C), chelerythrine chloride (D-F), harmane (G-I), and 18-crown-6 ether (J-L) reveal the robustness of the resistance and susceptibility phenotypes displayed by SAsmr, PAsmr, and FTsmr. The concentration and SMR being tested are listed in each graph.

**Figure S6:**
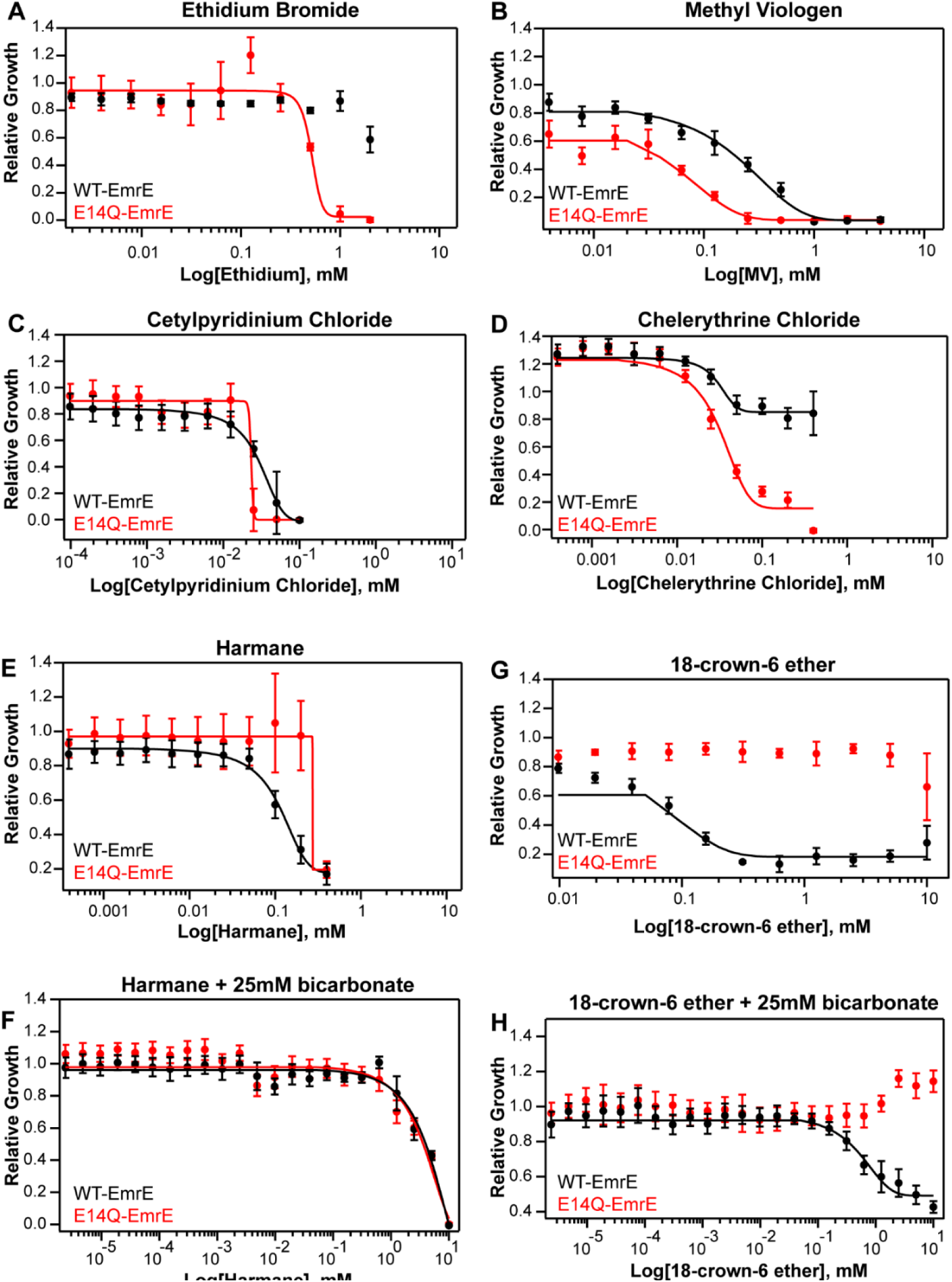
Dose-response curves of EmrE. Dose-response curves of MG1655 Δ*emrE E. coli* expressing WT-EmrE (black) or non-functional, E14Q-EmrE (red) in the presence of 6 substrates (A-D,E,G), and susceptibility substrates + 25mM bicarbonate (F, H). Each data point is the average of nine biological and technical replicates and error bars represent the standard deviation of all replicates. Curves were fit to a basic sigmoid curve using IGOR Pro Version 8. Curve fitting was not done when inhibition was not observed.

**Figure S7:**
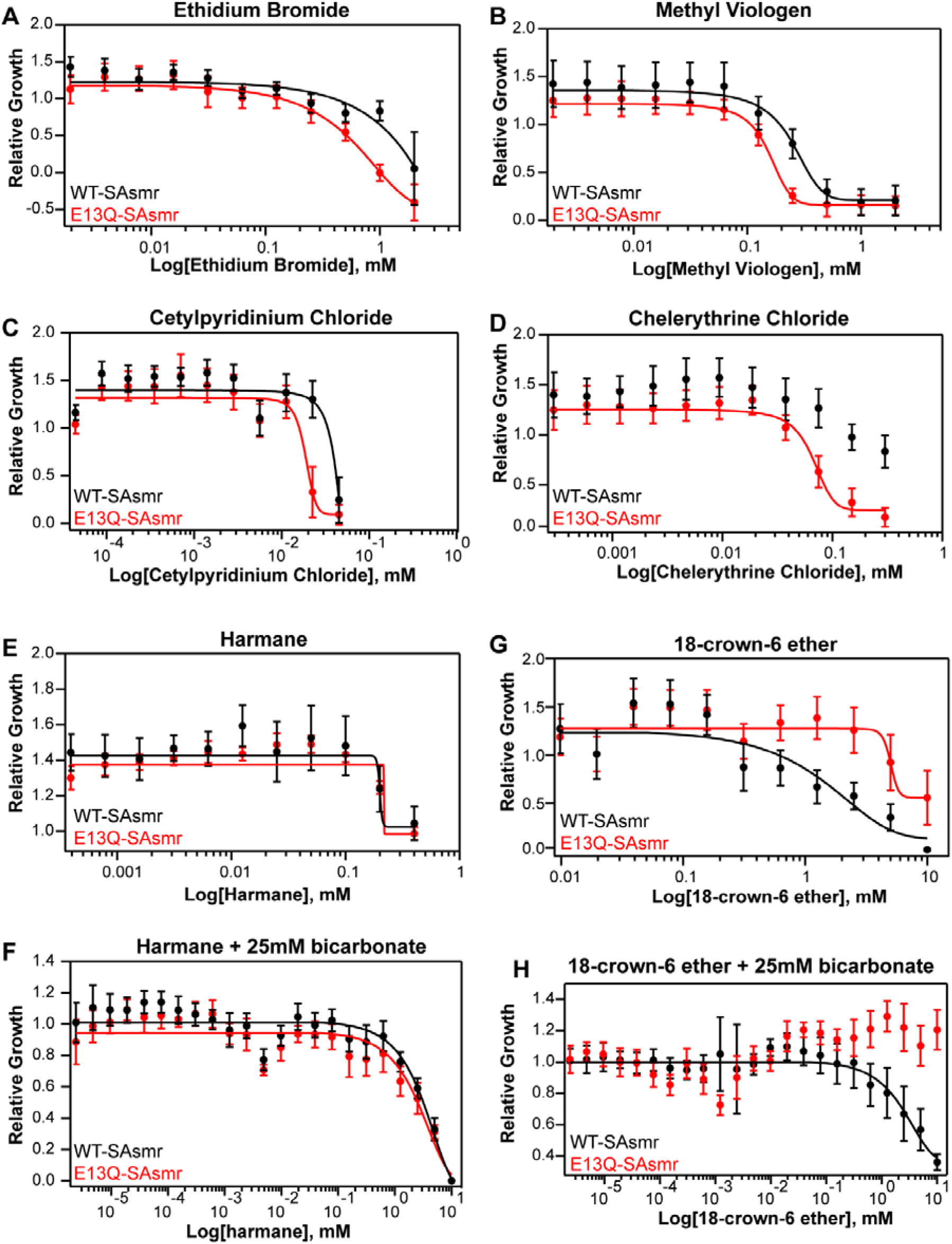
Dose-response curves of SAsmr. Dose-response curves of MG1655 Δ*emrE E. coli* expressing WT-SAsmr (black) or non-functional, E14Q-SAsmr (red) in the presence of 6 substrates (A-D,E,G), and susceptibility substrates + 25mM bicarbonate (F, H). Each data point is the average of nine biological and technical replicates and error bars represent the standard deviation of all replicates. Curves were fit to a basic sigmoid curve using IGOR Pro Version 8. Curve fitting was not done in the absence of inhibition.

**Figure S8:**
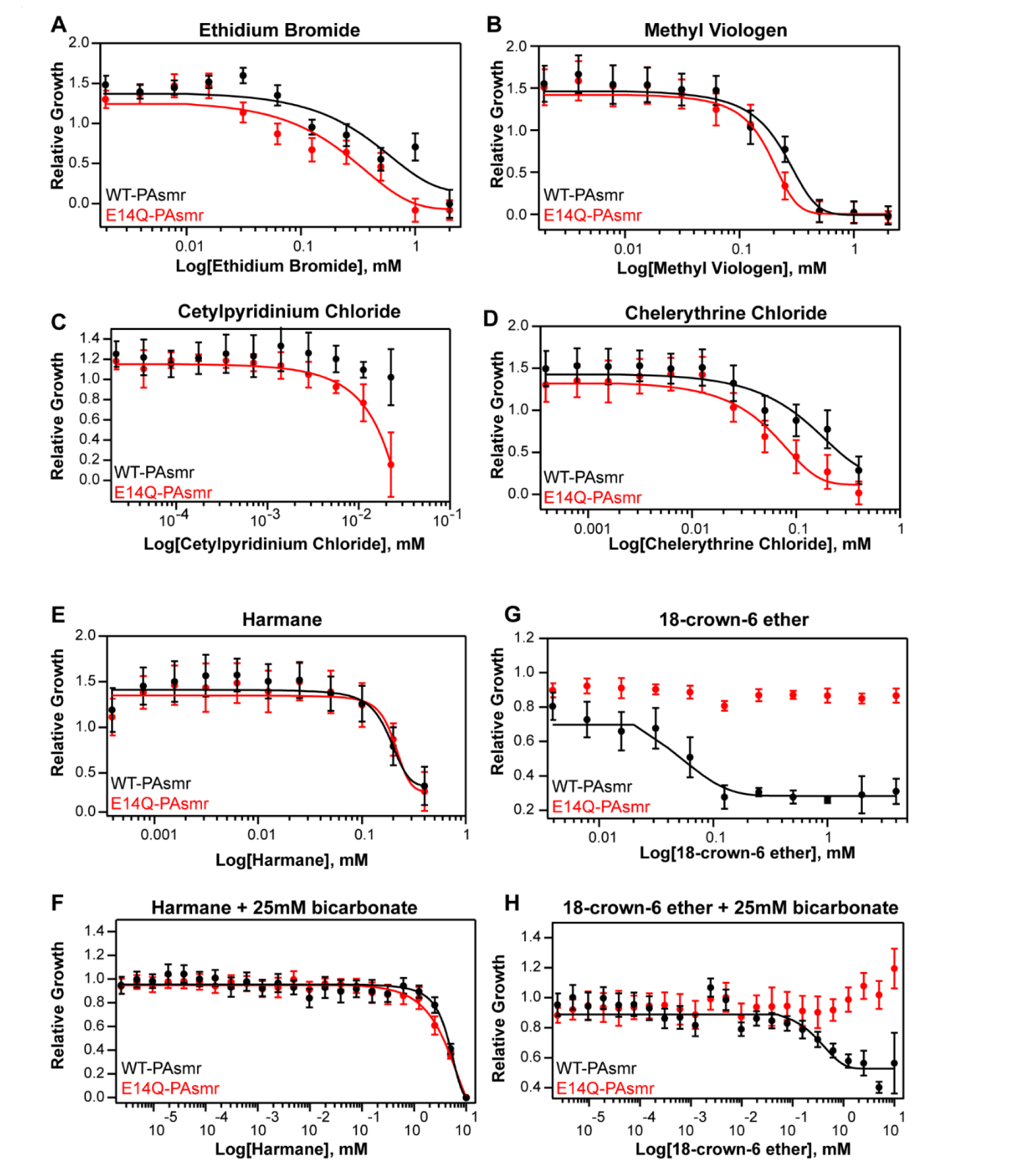
Dose-response curves of PAsmr. Dose-response curves of MG1655 Δ*emrE E. coli* expressing WT-PAsmr (black) or non-functional, E14Q-PAsmr (red) in the presence of 6 substrates (A-D,E,G), and susceptibility substrates + 25mM bicarbonate (F, H). Each data point is the average of nine biological and technical replicates and error bars represent the standard deviation of all replicates. Curves were fit to a basic sigmoid curve using IGOR Pro Version 8. Curve fitting was not done in the absence of inhibition.

**Figure S9:**
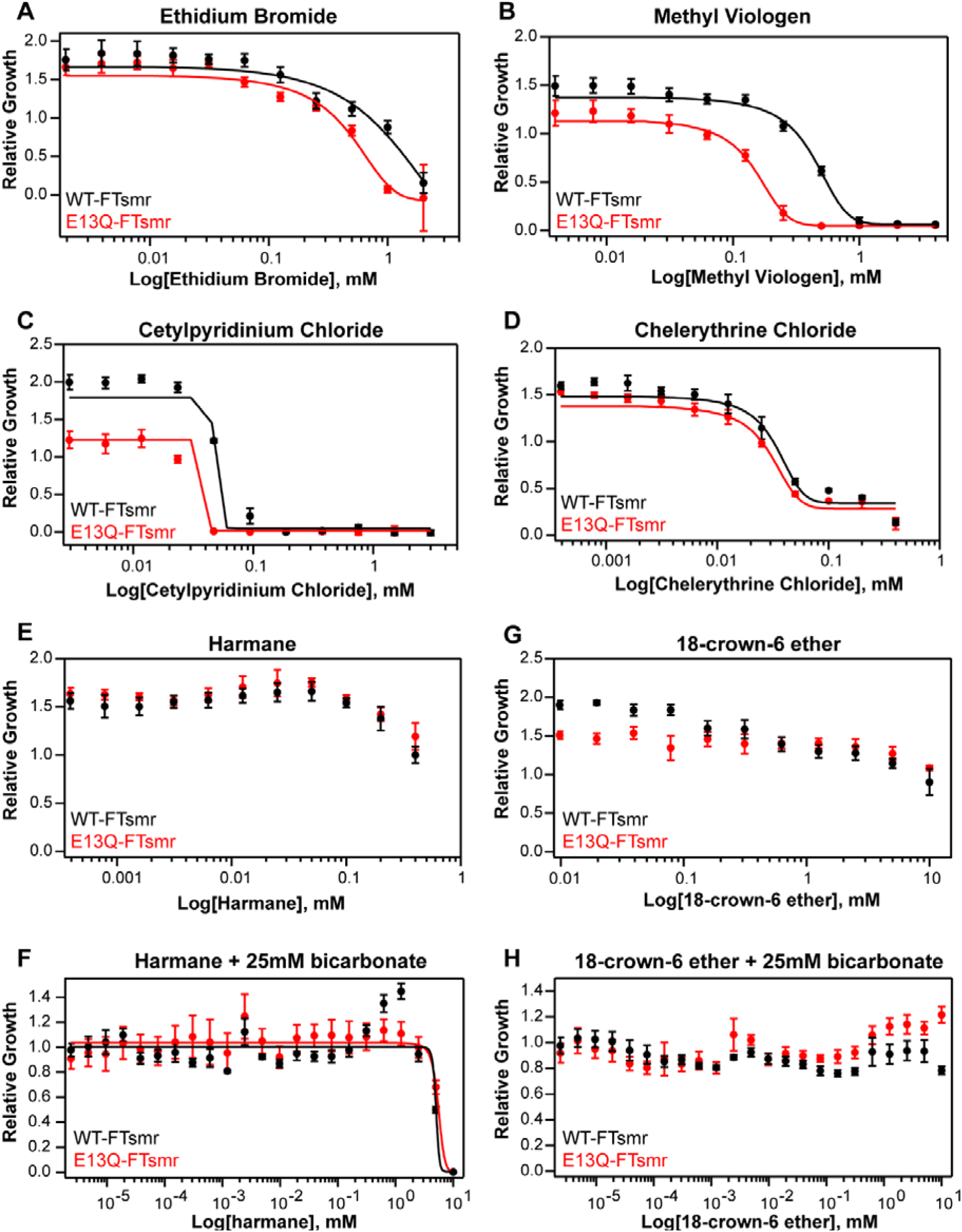
Dose-response curves of FTsmr. Dose-response curves of MG1655 Δ*emrE E. coli* expressing WT-FTsmr (black) or non-functional, E14Q-FTsmr (red) in the presence of 6 substrates (A-D,E,G), and susceptibility substrates + 25mM bicarbonate (F, H). Each data point is the average of nine biological and technical replicates and error bars represent the standard deviation of all replicates. Curves were fit to a basic sigmoid curve using IGOR Pro Version 8. Curve fitting was not done in the absence of inhibition.

**Figure S10:**
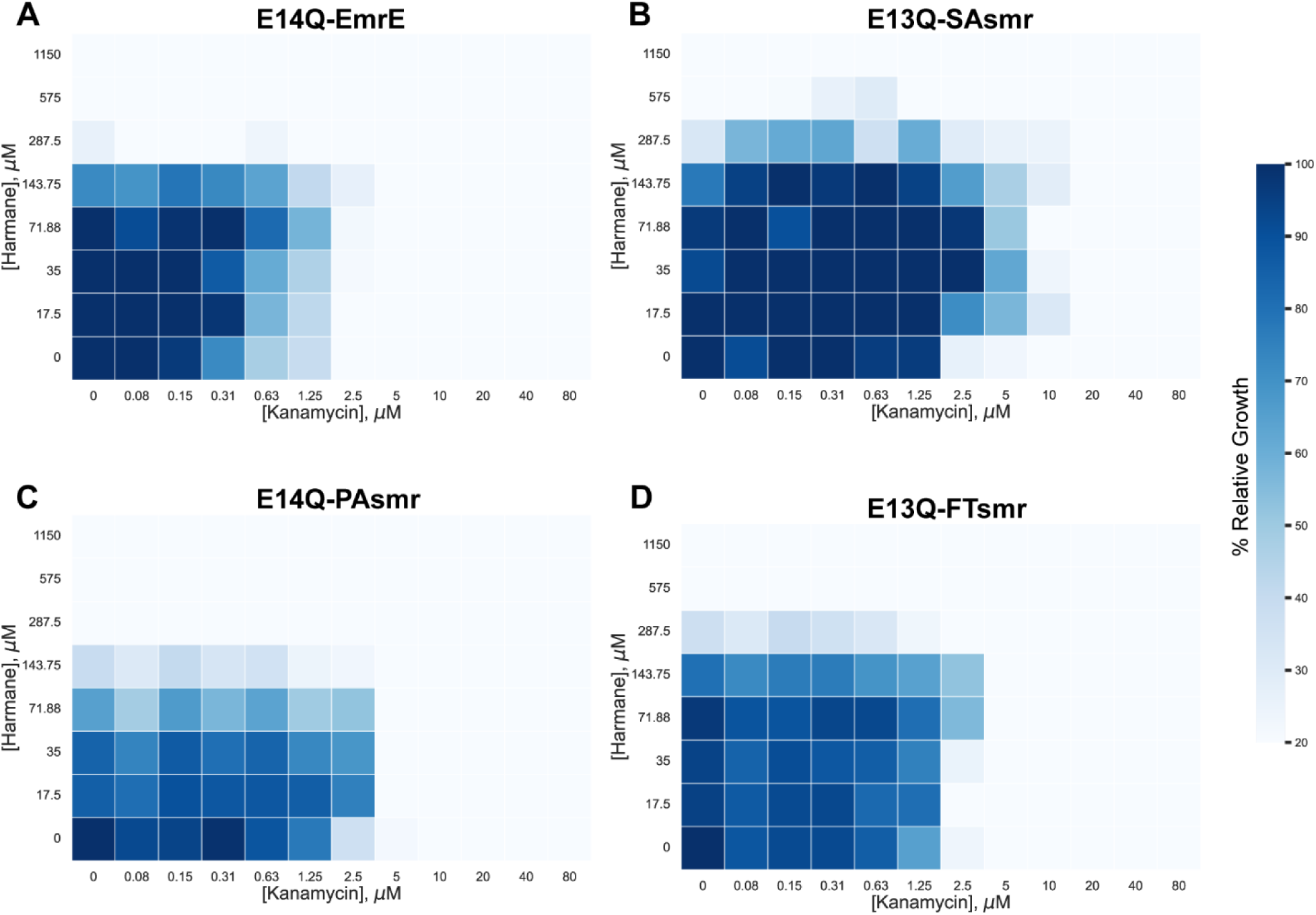
Harmane does not act synergistically with kanamycin in the presence of non-functional SMRs. Checkerboard assays of MG1655 Δ*emrE E. coli* cells expressing non-functional EmrE (A), SAsmr (B), PAsmr (C), or FTsmr (D) indicate that functional SMR transporters are required to observe synergy between harmane and kanamycin. Each cell represents the average relative growth across three technical replicates. The data is colored by % relative growth compared to no drug (lower left well) as measured by OD_600_, with relative growth at or below 0.2 (80% inhibition) is shown in white.

**Figure S11:**
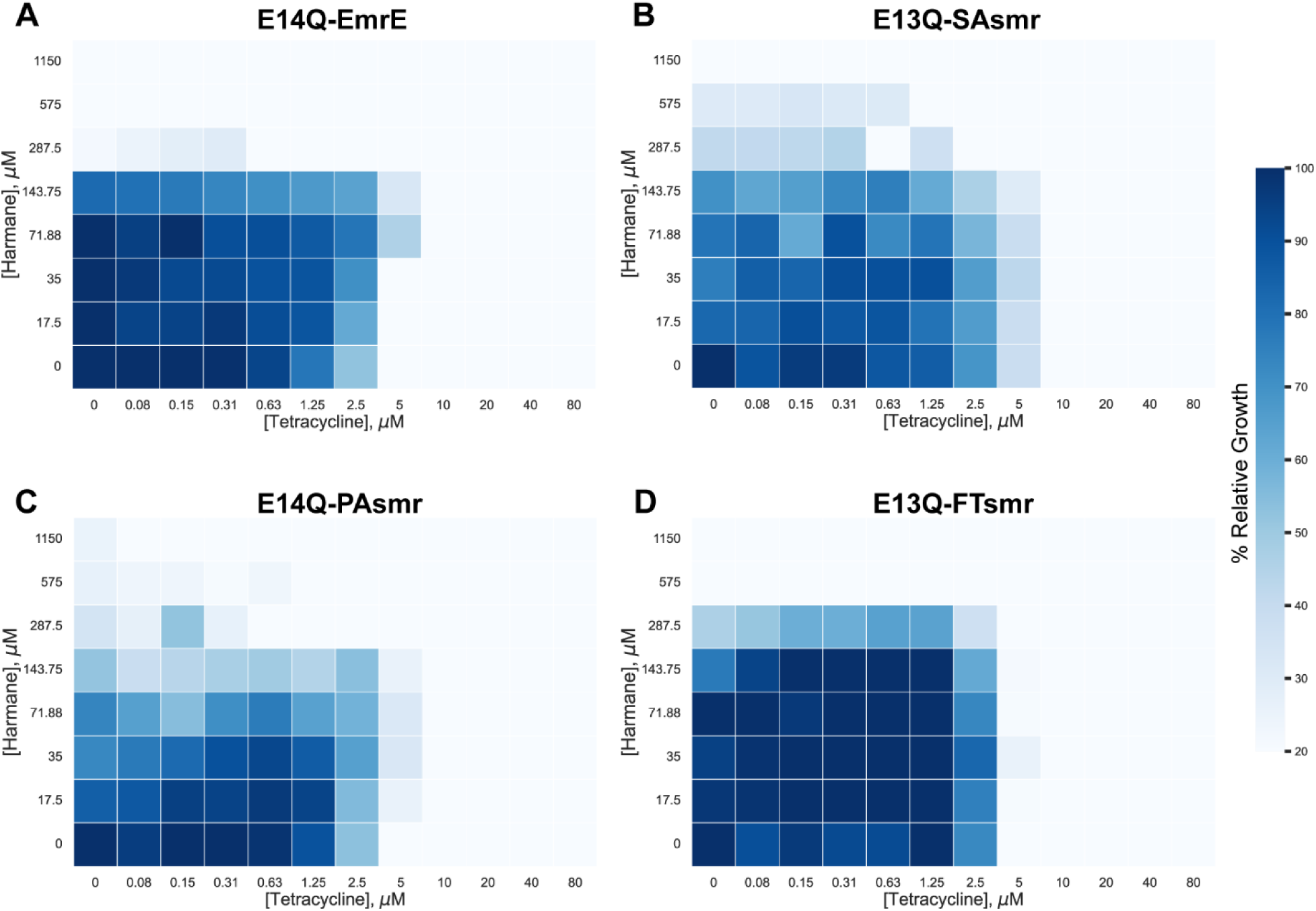
Harmane does not interact synergistically with tetracycline in the presence of non-functional SMR transporters. Checkerboard assays of MG1655 Δ*emrE E. coli* cells expressing non-functional EmrE (A), SAsmr (B), PAsmr (C), or FTsmr (D) demonstrate that there is no interaction between harmane and tetracycline in the absence of functional SMR transporters. Each cell represents the average relative growth across three technical replicates. The data is colored by % relative growth compared to no drug (lower left well) as measured by OD_600_, with relative growth at or below 0.2 (80% inhibition) is shown in white.

**Table S1:**
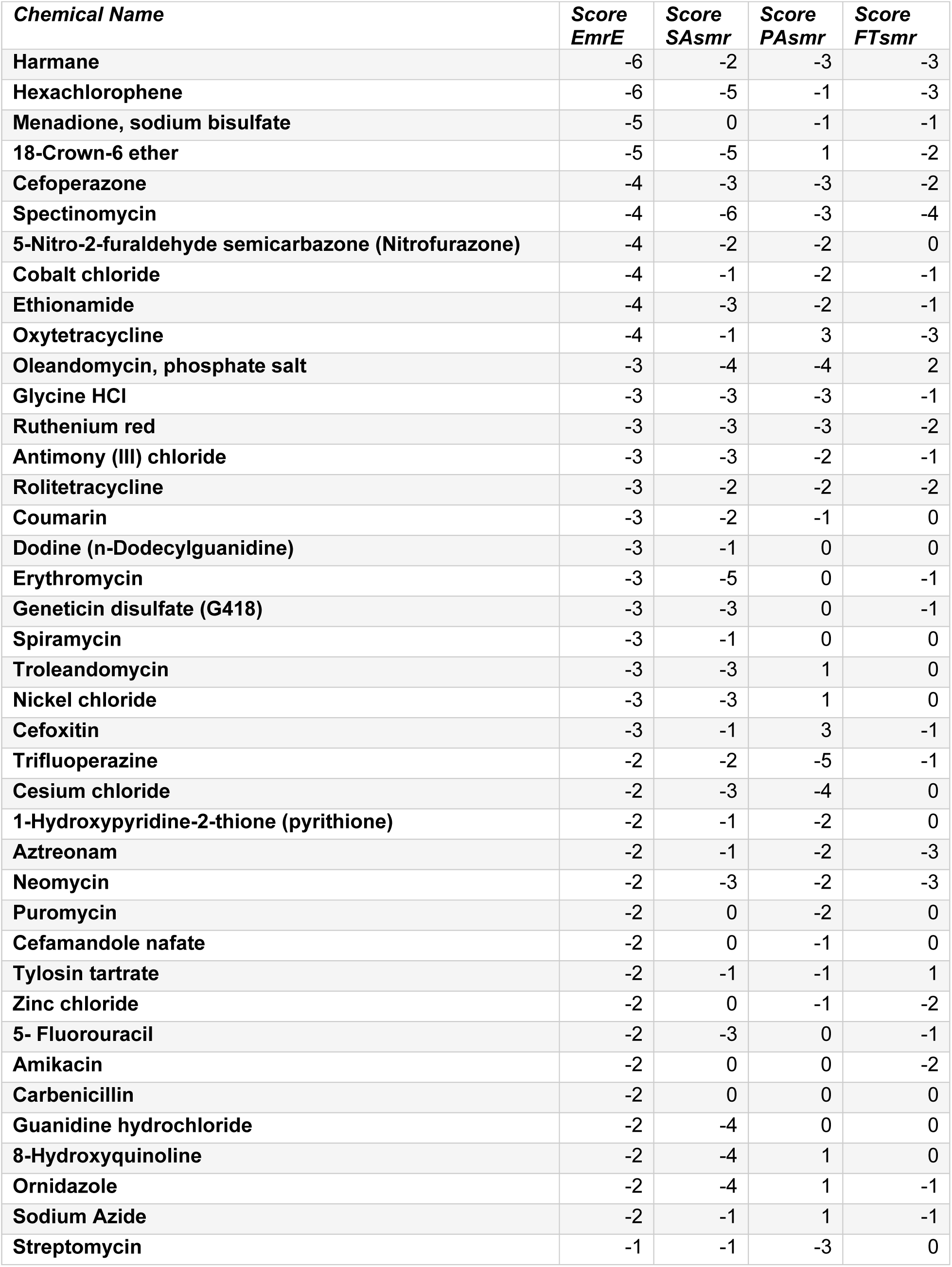
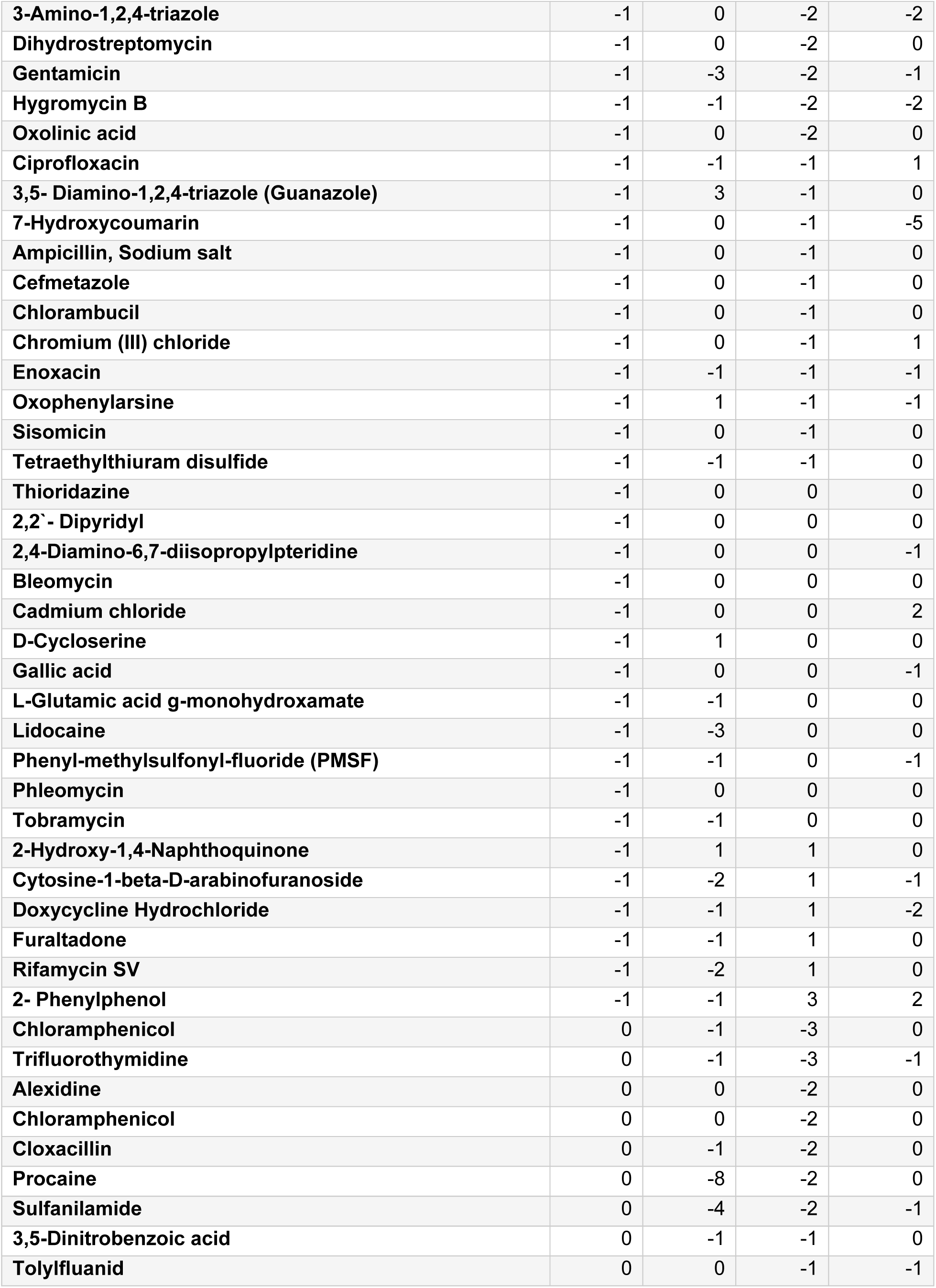
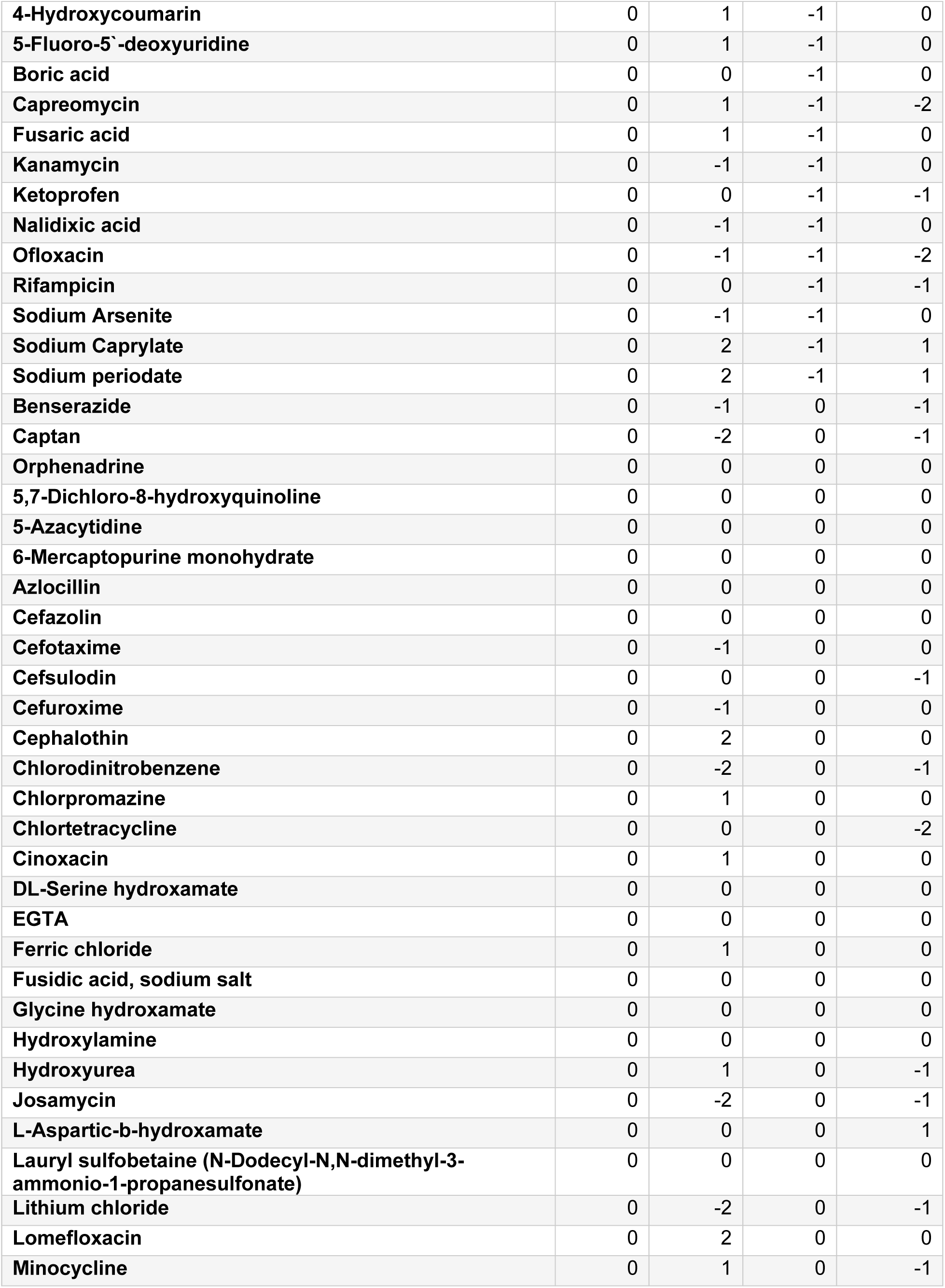
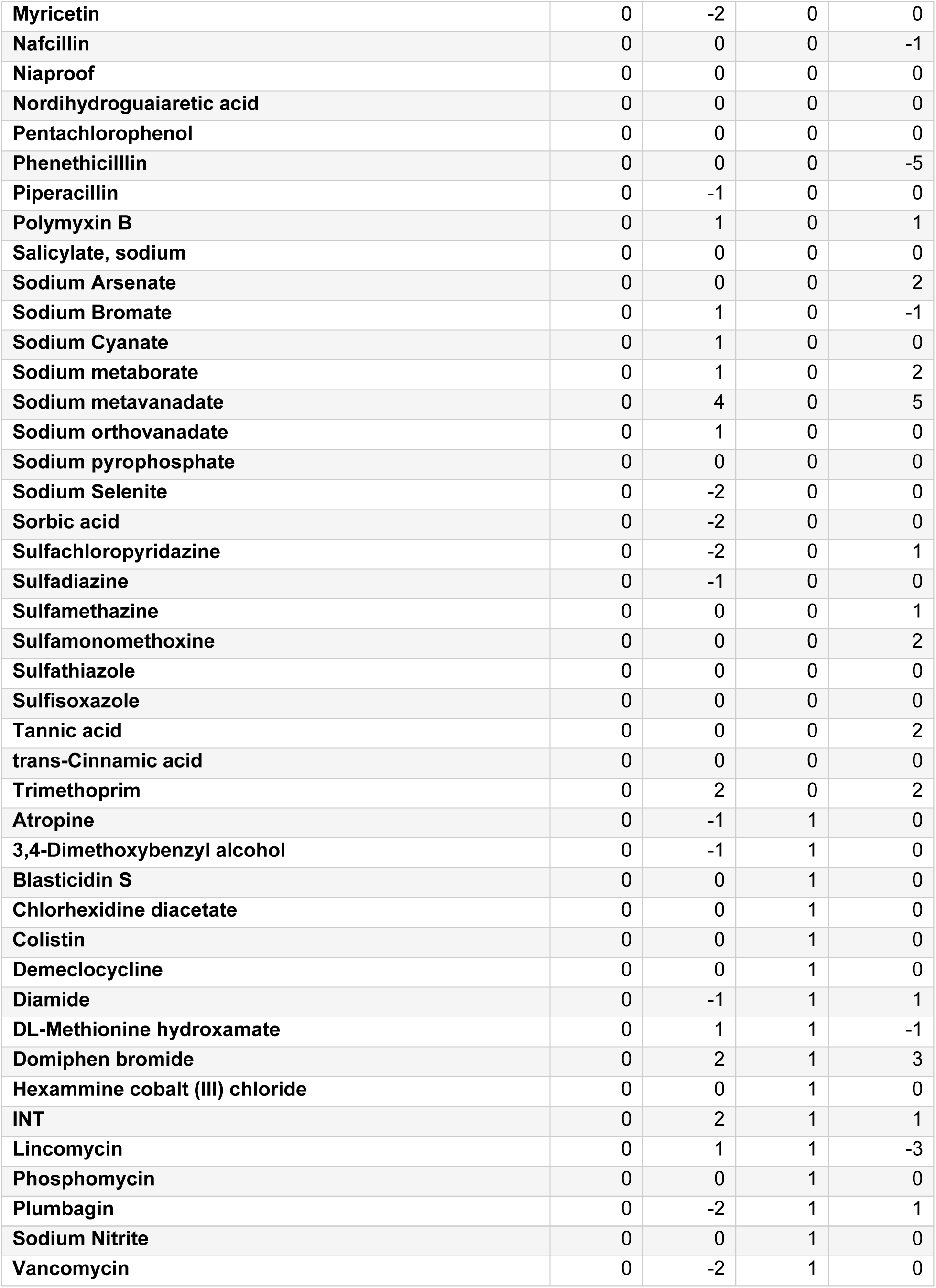
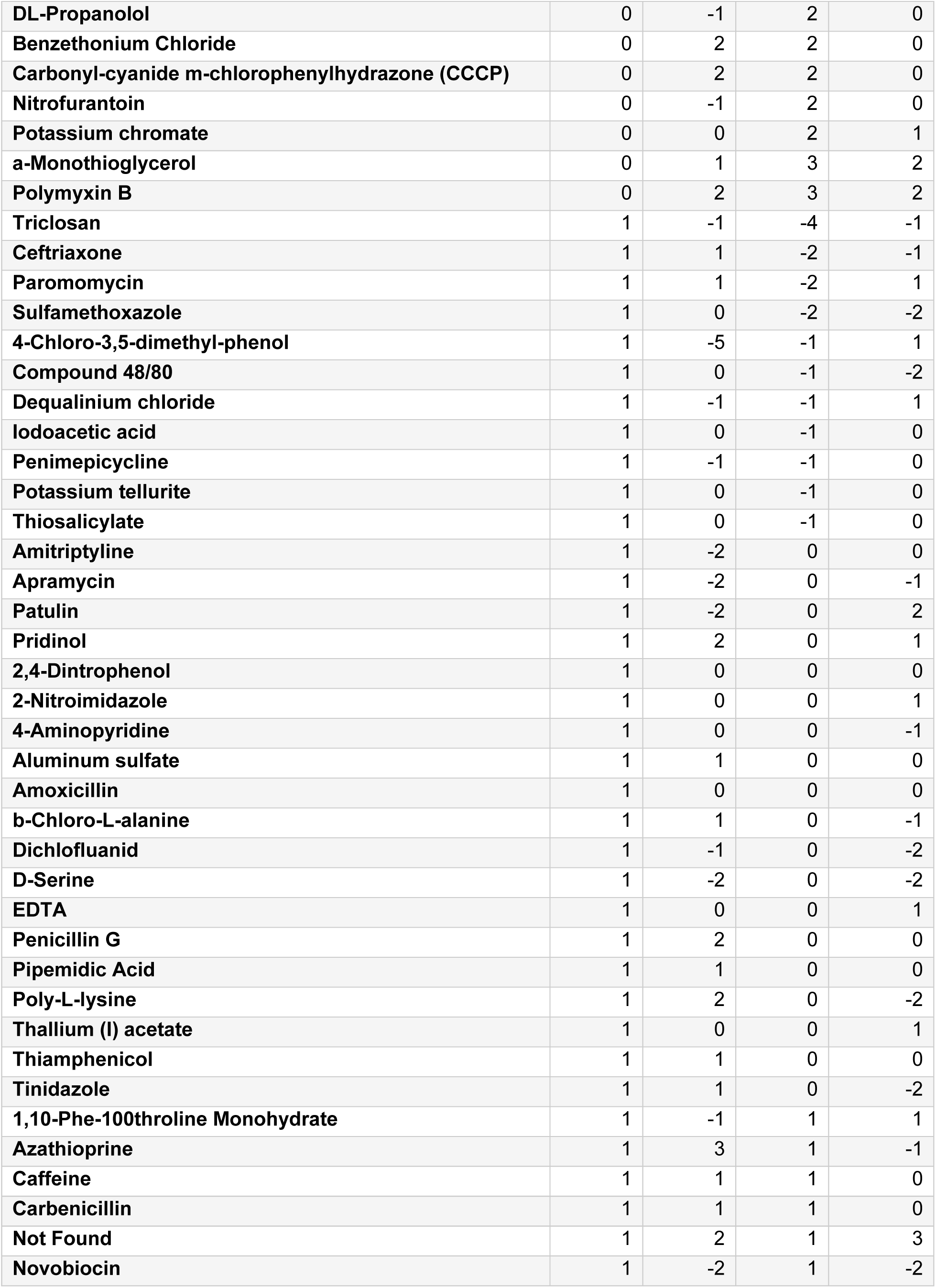
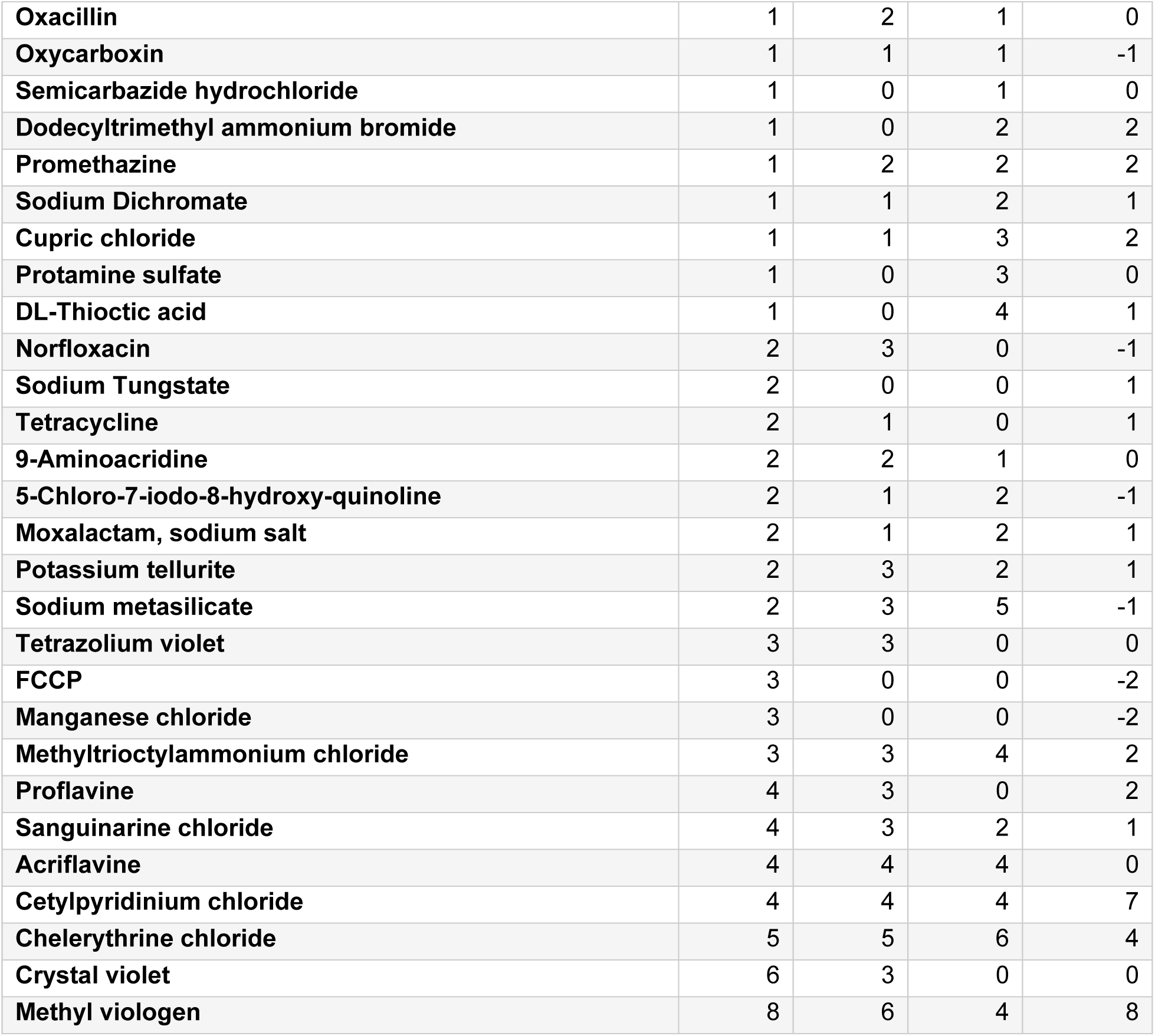
Biolog Hits sorted by EmrE Hit Score

